# Large-scale classification of metagenomic samples: a comparative analysis of classical machine learning techniques vs a novel brain-inspired hyperdimensional computing approach

**DOI:** 10.1101/2025.07.06.663394

**Authors:** Jayadev Joshi, Fabio Cumbo, Daniel Blankenberg

**Author notes:** These authors contributed equally to this paper (Jayadev Joshi, Fabio Cumbo). To whom correspondence should be addressed: Daniel Blankenberg, 9500 Euclid Avenue, NA29, Cleveland, OH 44195, USA.

## Abstract

Classical machine learning techniques have revolutionized bioinformatics, enabling researchers to extract knowledge from complex biological data. However, these techniques often struggle with high-dimensional data, where the increasing number of features leads to decreased performance, also affecting models accuracy. To address this problem, we explore hyperdimensional computing (HDC), an emerging brain-inspired computational paradigm that leverages high-dimensional vectors and simple arithmetic operations to represent and manipulate complex patterns, as an alternative approach in the context of supervised machine learning. In this work, we present a comprehensive comparative analysis of HDC against established machine learning techniques across a range of classification tasks. As a representative use case, we focus on classifying heterogeneous metagenomic samples based on their quantitative microbial profiles, using publicly available microbiome datasets. Our results demonstrate that HDC achieves comparable, and in some cases, superior classification accuracy to classical methods. Furthermore, our findings highlight the potential of HDC for improved computational efficiency, particularly when dealing with large-scale datasets, suggesting the HDC-based classifier as a promising tool for bioinformatics research, particularly in areas characterized by high-dimensional data. We also offer a Galaxy powered toolset to analyze your own datasets and generate reproducible workflows and adopt these methods in your own research with ease. Our investigation into the application of a HDC-based supervised machine learning technique for classifying microbial profiles in metagenomic samples yielded promising results, demonstrating the potential of this novel computational paradigm to complement and, in some cases, surpass the performances of well established machine learning techniques.

**Importance:** The growing complexity and dimensionality of biological data require more efficient and scalable machine learning approaches. HDC offers a novel alternative to conventional methods, showing resilience to high-dimensionality while maintaining competitive accuracy. This study demonstrates the effectiveness of HDC in classifying metagenomic samples based on their microbial composition. Our results suggest that HDC not only matches, but sometimes exceeds the performance of well-established methods. We make this approach accessible to the broader bioinformatics community with an open-source tool fully integrated into the Galaxy platform, facilitating its adoption and reproducibility, with the aim of integrating HDC into mainstream biological data analysis pipelines, especially for complex, high-dimensional tasks in microbiome research.

## Introduction

Metagenomics, the study of the whole genetic material recovered in a biological sample, has revolutionized our understanding of microbial communities and their impact on diverse ecosystems, from the human gut to the depths of the ocean [1]. Unlike traditional culture-based methods, computational metagenomics analyses provide a comprehensive in-silico view of microbial diversity and function, enabling the identification and characterization of both culturable and unculturable microorganisms [2]. A key aspect of metagenomics analysis involves quantifying the abundance of different microbial taxa within a sample, often referred to as microbial profiling [3]. This quantitative information provides insights into the composition of microbial communities and how they vary across different environments or experimental conditions [4].

High-throughput sequencing technologies have enabled the generation of vast amounts of metagenomic data, leading to the availability of public pre-computed datasets of quantitative microbial profiles [5–9]. These profiles, typically represented as high-dimensional vectors, capture the absolute and relative abundance of thousands of microbes within each sample at different taxonomic levels [10]. Analyzing such datasets requires sophisticated approaches, with supervised machine learning emerging as a powerful tool for extracting significant insights [11]. Supervised learning algorithms, trained on labeled datasets, can learn to predict specific outcomes or classify samples based on these microbial profiles [12].

Classical machine learning algorithms, like decision trees, random forest, support vector machine, and logistic regression, have been successfully applied to a wide range of classification tasks in omics research, including those involving microbes [4,5,13,14]. These include predicting host phenotypes based on the microbiome composition, identifying disease biomarkers from microbial profiles, and classifying environmental samples based on their microbial communities [15]. However, the inherent high dimensionality of microbial profiles presents challenges for these classical methods [16]. As the number of features, i.e., microbial taxa, increases, especially now that we are finally able to detect still unknown microbes in metagenomic samples, the performance of these algorithms can deteriorate, affecting models accuracy and leading to misclassifications [17,18].

To address these challenges, we explore hyperdimensional computing (HDC), an emerging brain-inspired computational paradigm [18]. It leverages high-dimensional vectors to represent data and uses simple arithmetic operations to manipulate and represent complex relationships between them. It already found applications in a wide series of fields, including bioinformatics, natural language processing, machine learning, AI, and other scientific domains [18–22]. Here we investigate how a HDC-based supervised classification technique could bring any advantage in terms of performances and models accuracy compared to classical machine learning approaches in the context of discriminating heterogeneous metagenomics samples based on their quantitative microbial profiles.

## Materials and Methods

Here we present a detailed overview of the datasets involved in our study, following the mathematical model underlying our supervised classification method based on the HDC paradigm. We also present a brief overview of the classical machine learning techniques and the evaluation metrics considered in our comparative analysis.

### Data Sources

Our study focuses on public datasets available under the *curatedMetagenomicData* [10] package for R, which offers standardized, curated human microbiome data, including gene families, marker abundance, marker presence, pathway abundance and coverage, and relative abundances. All these datasets were collected from approximately 168 different public experimental studies. Relative abundance profiles for each of the samples in this package have been computed with *MetaPhlAn3* [23], a marker-based profiler able to detect and estimate the abundance of bacteria, archaea, and fungi in metagenomic samples. Alongside this data, the R package also provides a set of manually curated and standardized metadata describing samples, the experimental conditions at the time of sequencing, and other demographic and health information about donors.

In our study, we consider 38,000 samples divided across various categories, which include birth method, history of periodontitis, disease subtype, study condition, birth order, country, breastfeeding practice, non-westernized, age category, family role, location, gender, body site, birth control pill, smoker, diet type, and dental sample type. Initially, the dataset includes a total of 25 categories, and each category combines samples from different studies. We discard data from specific studies where the number of samples is less than 100. We also compute a normalized score to determine the balance between binary class data instances. Figure 1’s box plot depicts the data balance across binary classes, where a score of 1 indicates perfect balance with both classes having 50% of the samples, while a score close to 0 indicates highly imbalanced data. The score is calculated based on the class containing fewer samples divided by the total number of samples in the dataset. This score is then normalized using the min-max normalization based on the score calculated across all datasets.

**Figure 1.**
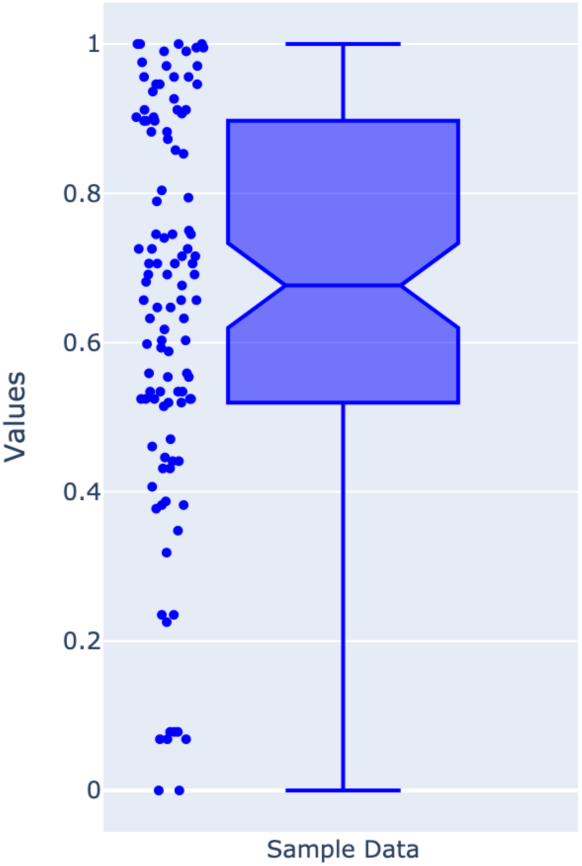
Box plot representing the data distribution across the two classes. Datasets that show a score close to 1 indicate balanced data, while datasets with a score close to 0 indicate highly imbalanced data.

The selected studies’ scores range from 1.0 to 0.6, indicating the datasets range from perfectly balanced (1.0) to moderately balanced (0.6), making them suitable for a classification task. A detailed sample distribution across two classes of selected studies were represented in Figures S1-5 in Supplementary Material. Studies which never fall under this range are discarded from this study. Additionally, Table 1 reports the set of categories considered for stratifying the datasets with the number of datasets involving those categories and their long description.

**Table 1:** This table shows the list of considered metadata categories alongside the number of datasets that involve those categories, and their long description retrieved from the *curatedMetagenomicData* package for R.

| Category | Datasets | Long Description |
| --- | --- | --- |
| *Age category* | 9 | The category of the age based on different thresholds. Possible values: *newborn* (age $\leq 1$), *child* (age $\leq 12$), *adult* ($18 \leq \text{age} \leq 65$), *senior* (age $> 65$) |
| *Birth control pill* | 1 | Whether a donor takes a birth control pill at the time of collecting the metagenomic sample. Possible values: *yes*, *no* |
| *Born-method* | 2 | The birth delivery method. Possible values: *C section*, *vaginal* |
| *Birth order* | 1 | In the case of twins, who is born first. Possible value: 1, 2 |
| *Body site* | 1 | The body site where a metagenomic sample has been collected from. Possible value: *stool*, *oral cavity* |
| *Breastfeeding practice* | 1 | The practice used to breastfeed a newborn. Possible values: *exclusively breastfeeding*, *mixed feeding* |
| *Country* | 3 | Geographic location where the donor of a metagenomic sample comes from. Possible values: *El Salvador*, *Peru*, *Germany*, *Kazakhstan*, *Denmark*, *Spain* |
| *Dental sample type* | 1 | Type of metagenomic dental sample. Possible values: *implant*, *teeth* |
| *Disease subtype* | 2 | Whether a donor is affected by a specific disease, this is used to specify the subtype of the disease. Possible values: *ulcerative colitis*, *Crohn's disease* |
| *Family role* | 2 | The role of a donor in their family. Possible values: *mother, child* |
| *Gender* | 33 | Donor's gender. Possible values: *male, female* |
| *Diet Type* | 1 | Whether a donor follows a gluten-based diet. Possible values: *high gluten diet, low gluten diet* |
| *History of periodontitis* | 1 | Whether a donor has a medical history of periodontitis. Possible values: *yes, no* |
| *Location* | 3 | The specific place where the donor of a metagenomic sample used to live. Possible values: *rural village, Lima periurban shantytown, Kerala, Bhopal, Boston, Atlanta* |
| *Non-Westernized* | 1 | Whether a donor's lifestyle is close to a non- EU, USA, and Canadian culture. Possible values: *yes, no* |
| *Smoker* | 3 | Whether a donor is a smoker at the time of collecting the metagenomic sample. Possible values: *yes, no* |
| *Study condition* | 10 | Whether a donor is affected by a specific condition at the time of collecting a sample. Possible values: *control, T2D, ME/CFS, STH, CRC, ACVD, cirrhosis, schizophrenia, IBD* |
| ORR<br>(Objective Response Rate) | 1 | Objective Response Rate (ORR) is a clinical trial metric used to measure the proportion of patients whose cancer shrinks or disappears after treatment. Possible values: *yes, no* |

### Hyperdimensional Computing and the MAP model

The HDC paradigm takes inspiration from how the human brain represents and manipulates information. It consists in encoding every kind of data into random vectors in a high-dimensional space, called hypervectors, and combining them using a small set of simple arithmetic operations to represent complex relationships between data. Here we focused on the MAP model, an acronym that stands for Multiply-Add-Permute, that limits the arithmetics to three mathematical operations on vectors, i.e., the element-wise multiplication (a.k.a. binding), the element-wise addition (a.k.a. bundling), and the permutation which consists of rotating a vector by a fixed number of positions.

These three simple operations brings different properties when applied to manipulate and combine vectors:

- <u>Bundling (element-wise addition)</u>: this operation, defined as *C* = *A* + *B*, creates a superposition of the input vectors. The resulting vector *C* is an average that contains information from both *A* and *B*. Consequently, the output vector *C* will have a high cosine similarity to both of its input vectors. This property is used to group items into a set or accumulate evidence, as the resulting set vector remains similar to its constituent items;
- <u>Binding (element-wise multiplication)</u>: this operation, defined as *C* = *A* × *B*, is used to form a new, distinct association between two vectors. The resulting vector *C* is quasi-orthogonal (dissimilar) to both *A* and *B*. Binding is also a similarity-preserving transformation: if two vectors *A* and *Ā* are highly similar, their bound products with a third vector *X* (i.e., *A* × *X* and *Ā* × *X*) will also be highly similar to each other. This property is critical for robust matching. The operation is also invertible (e.g., (*A* × *B*) × *B* = *A* in bipolar systems) and distributes over bundling;
- <u>Permutation (rotation)</u>: this operation, *C* = ρ(*A*), deterministically shuffles the elements of *A*. Like binding, permutation produces a new vector *C* that is quasi-orthogonal (dissimilar) to the input vector *A*. This is essential for encoding order and position. Permutation is also a similarity-preserving transformation, meaning if *A* and *B* are similar, ρ(*A*) and ρ(*B*) will be similar to each other by the same degree. It is invertible and distributes over both bundling and binding.

Here we are going to provide some additional insights about how to encode information as hypervectors and how to build a supervised classifier according to the MAP model. In order to do so, we take inspiration from the *chopin2* [24] software package originally designed to discriminate between human healthy and tumor samples based on their DNA-Methylation profiles characterized by over 450-thousand features (i.e., CpG islands). The authors of *chopin2* show great classification results and models with high accuracy rate able to run on commodity hardware. Here we show our interpretation of the supervised HDC-based classification model.

#### Encoding

According to the HDC paradigm, every piece of information must be encoded as a random binary (0, 1) or bipolar (-1, 1) vector in a high-dimensional space, typically around 10,000 dimensions. The high dimensionality is crucial because, in order to distinguish between two vectors representing different pieces of information, we rely on computing their distance or similarity (e.g., Hamming distance or cosine similarity). The higher the dimensionality, the higher the chances that two randomly generated vectors are orthogonal or quasi-orthogonal in the same space. In a 10-thousand dimensional space, we can define at most 10-thousand orthogonal vectors, which sets an upper bound on the number of different types of information we can encode.

In the context of a classification model, given a numerical matrix where rows represent samples and columns represent features, every numerical value must be encoded into a hypervector. Since the spectrum of all possible numbers is infinite, we start by establishing how many vectors we want to define *a priori* so that we can then map the numerical values in the matrix to the set of hypervectors. For instance, if we define 100 hypervectors and the numerical values in our dataset range from 0 to 100, we can then map all the numbers between 0 and 1 to the first hypervector, numbers between 1 and 2 to the second hypervector, and so on, up to the numbers between 99 and 100 that must be mapped to the last 100th hypervector.

However, all these 100 hypervectors are not random. We must be able to distinguish two different numbers in the same high-dimensional space, but we also want to maintain the information about the closeness of different numbers in the interval 0-100. More specifically, the number 0 is different from the number 1 and the number 100, but it is closer to the number 1 rather than the number 100, and this property must be also reflected in the vector space. In order to do so, we start building a random binary or bipolar vector which represents the base used to encode all the vector representations of the numerical values in our dataset. The numbers between 0 and 1 are mapped to this base vector, while the vector representing numerical values between 1 and 2 is built by flipping half of the bits of the base vector (0 becomes 1 in binary vectors, while -1 becomes 1 in bipolar vectors, and the other way around) in random positions. This process continues up to the definition of the very last hypervector representing numbers between 99 and 100 which is built by flipping half of the bits of the hypervector representing the previous numerical interval (i.e., 98-99 in this case) in random positions. Note that flipping half of the bits in a random vector is enough to make the original vector and the flipped one quasi-orthogonal in the space. The quasi-orthogonality of these vectors allows us to discriminate them in the same space when computing a distance or similarity measure, while the quasi- property of orthogonality also makes them closer to the vector representation of adjacent numerical intervals. These hypervectors are also called level vectors.

#### Training

The primary objective of the training phase is to build a model capable of accurately predicting the class label of a new sample. In order to build a model according to the HDC paradigm, we first have to encode every sample in the training set into a hypervector. We have already mapped every numerical value in the dataset to a level vector as we mentioned in the previous paragraph. Here we have to combine the vector representations of the numerical values under a specific row (sample) considering their position in the dataset, i.e., maintaining their order according to the order of the features, like in the following formula (1):

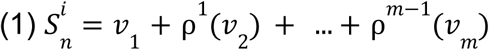

Here, the vector representation of a sample is defined as the element-wise sum (bundling) of all the level vectors representing the numerical values at its specific row in the dataset, permuted by their position in which they occur in the same row (i.e., the index of the features). Please note that permuting a vector is an invertible operation and preserves the similarity with the input vector, encoding, at the same time, the information about the position of that numerical value and, implicitly, which feature it belongs to.

Once we encode all the samples in the training set, we should collapse them all together according to their class. Collapsing means simply applying the element-wise sum (bundling), which produces a vector representation of a specific class that maintain, as a property of the bundling operation, a similarity with all the vectors it is composed of, as in the following formula (2):

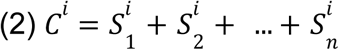

At the end of the training phase, we end up with a hypervector for each of the classes that group samples in our dataset. The set of classes hypervectors, together with the level vectors, represents the classification model. The complete procedure for generating this model is detailed in Algorithm 1.

##### Algorithm 1

This pseudocode details the training phase of the HDC model. The algorithm initializes a high-dimensional zero-vector for each class. It then iterates through every sample in the training set. For each individual sample, it constructs a unique sample hypervector (S_1) by sequentially processing each of its features. This involves mapping the numerical value of a feature to its corresponding level vector (v_level), applying a permutation (PERMUTE) based on the feature’s index (its position j), and bundling (element-wise addition) this permuted vector into the sample hypervector S_i. Once the full sample hypervector is built, it is bundled into the main hypervector representing its assigned class (Model[sample_label]). The final output is the Model, which consists of the set of class hypervectors that are then used for classification.

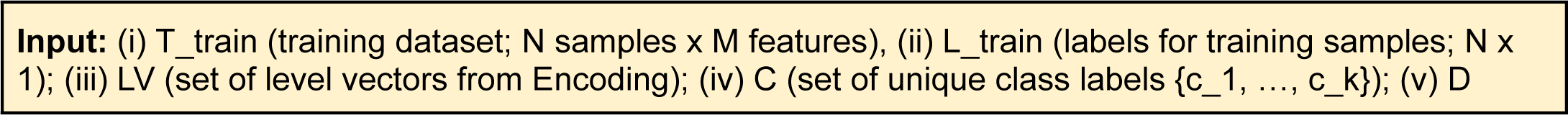
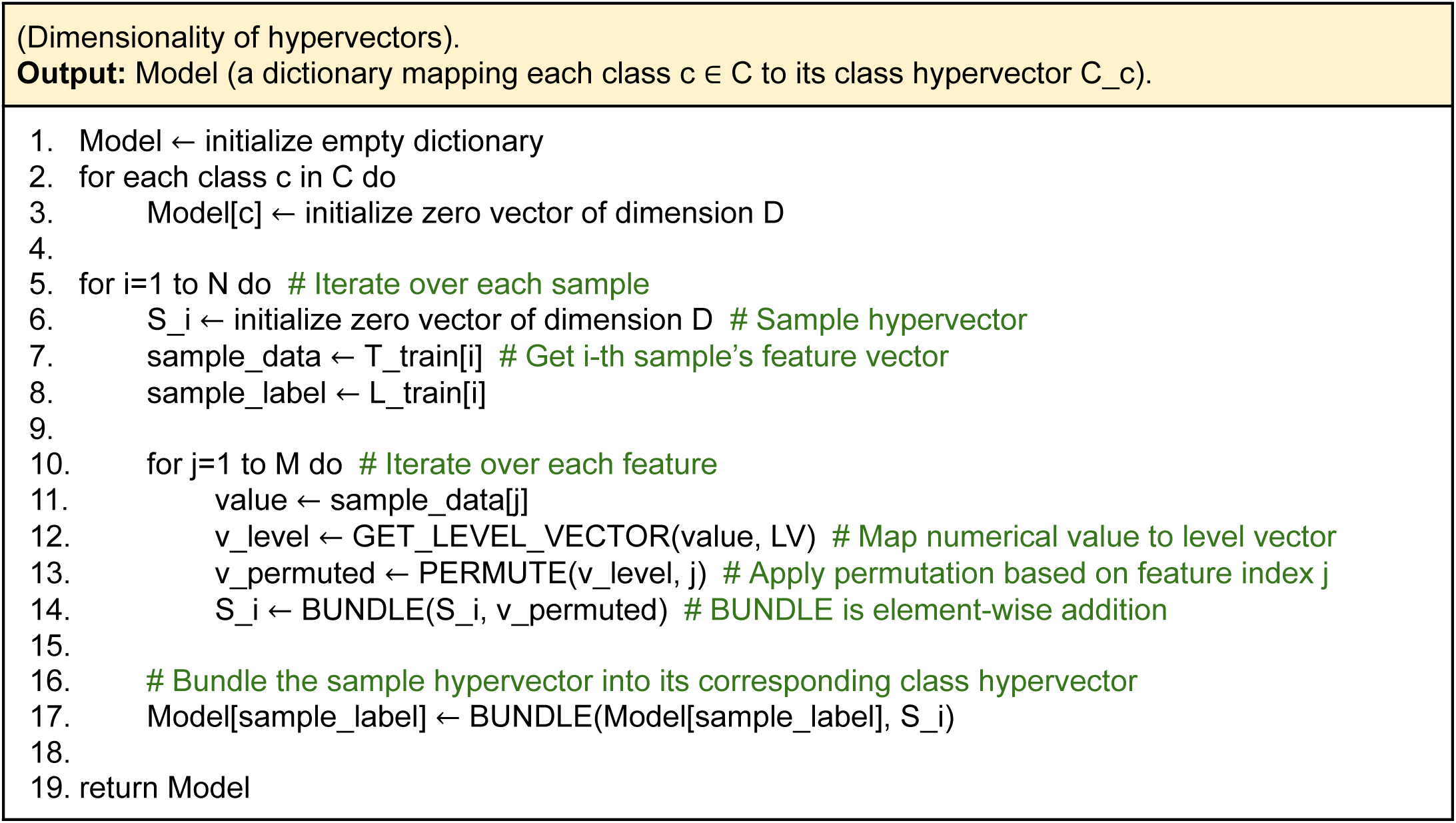

#### Testing

Testing a machine learning model means evaluating its performances in predicting the class of new samples that have not been previously seen during the training phase. Here, we first encode the samples in the training set in the same way we previously encoded the samples in the test set. Now, the prediction consists of computing the cosine similarity between each of the training sample hypervectors and the vector representation of the classes computed during the training phase. The predicted class is the closest one among the class vectors. The prediction process is formally described in Algorithm 2.

##### Algorithm 2

This pseudocode outlines the prediction phase. For each new sample in the test set, the algorithm first constructs its sample hypervector (S_test_i) using the same encoding process as in training (Algorithm 1). This involves mapping feature values to level vectors, permuting them by feature index, and bundling them together. Next, it computes the cosine similarity between the newly created test sample hypervector and every class hypervector stored in the trained Model. The sample is assigned the label of the class that yields the highest similarity score, as this represents the closest class in the high-dimensional space.

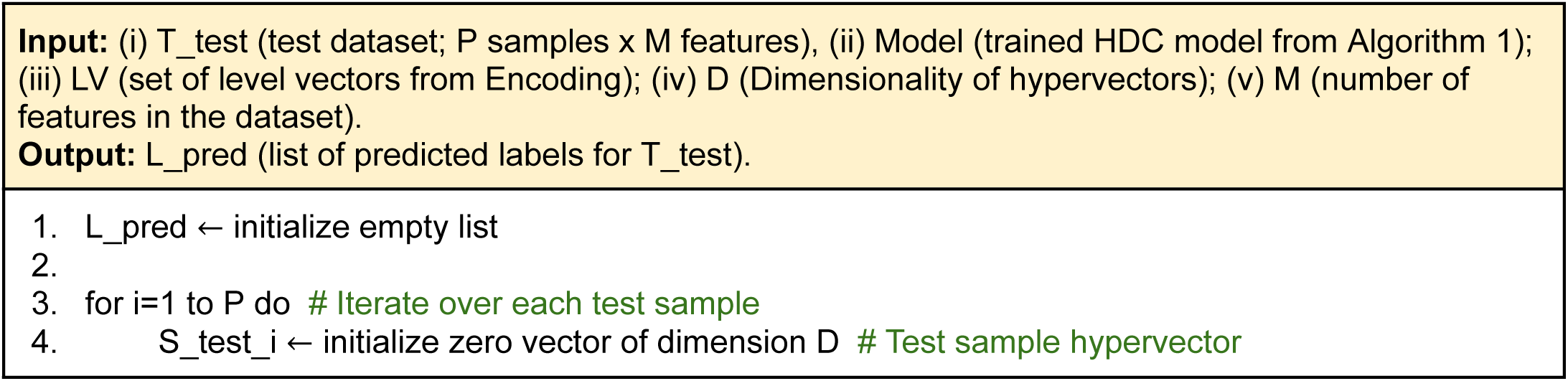
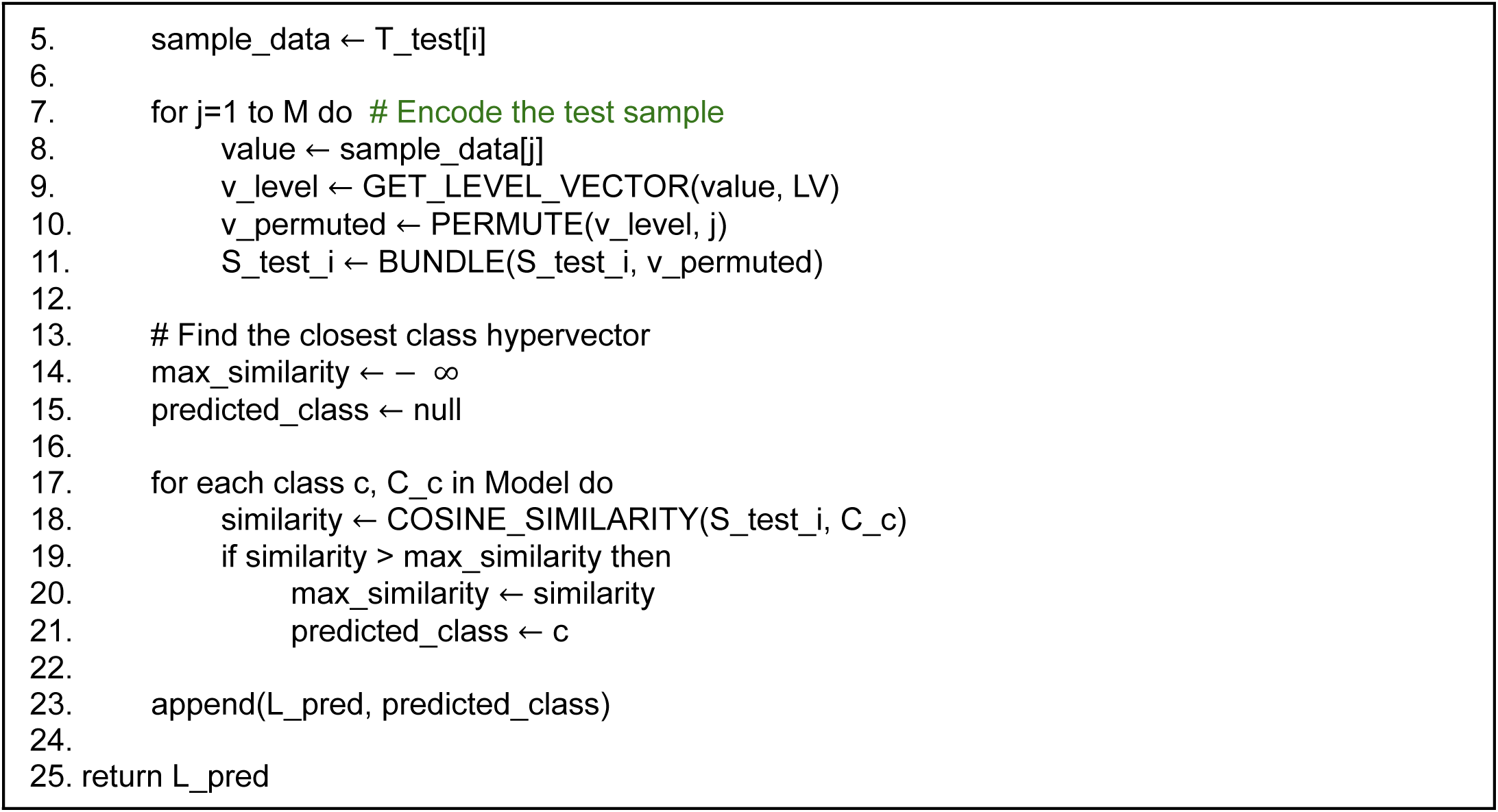

#### Mitigating the error-rate

As previously mentioned, the classification model consists of the class vectors that result from the bundling of all the vector representations of the training samples belonging to a specific class. The process of bundling two vectors preserves the similarity between the input vectors, but the more vectors we bundle, the more noise we introduce in our model. For this reason, we perform an iterative procedure called retraining with the aim of mitigating the model error rate.

We first define the error rate of a classification model as the number of misclassified training samples over the total number of training samples. If the error rate is greater than zero, we element-wise sum the vector representation of the misclassified training samples to their correct class vectors, and we element-wise subtract the same training vectors from their wrongly predicted class vectors, in order to increase and decrease their signal. This process runs iteratively until convergence, i.e., the error rate does not decrease anymore. This iterative retraining procedure is formalized in Algorithm 3.

##### Algorithm 3

This pseudocode details the iterative retraining process. The algorithm first pre-computes the sample hypervectors for all training samples. It then enters a loop, where each pass involves: (i) predicting the class for every training sample using the current model, (ii) identifying all misclassified samples, (iii) aggregating corrections by adding a misclassified sample’s hypervector to its correct class (BUNDLE) and subtracting it from its incorrectly predicted class (SUBTRACT). After checking all samples, the algorithm calculates the new error rate. If the error rate is zero or has not decreased since the last iteration, the loop terminates (convergence). Otherwise, the aggregated corrections are applied to the class hypervector, and the process repeats.

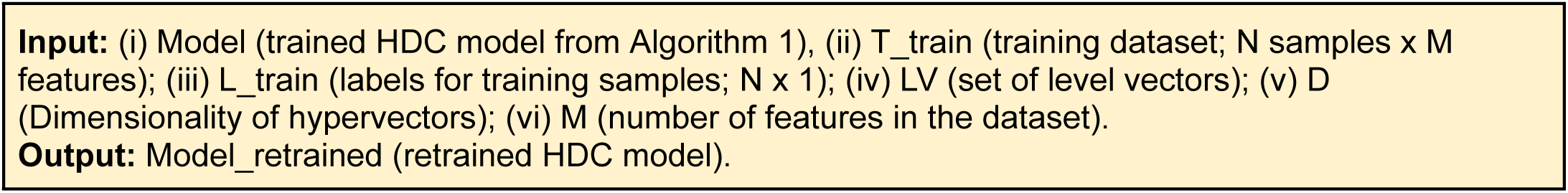
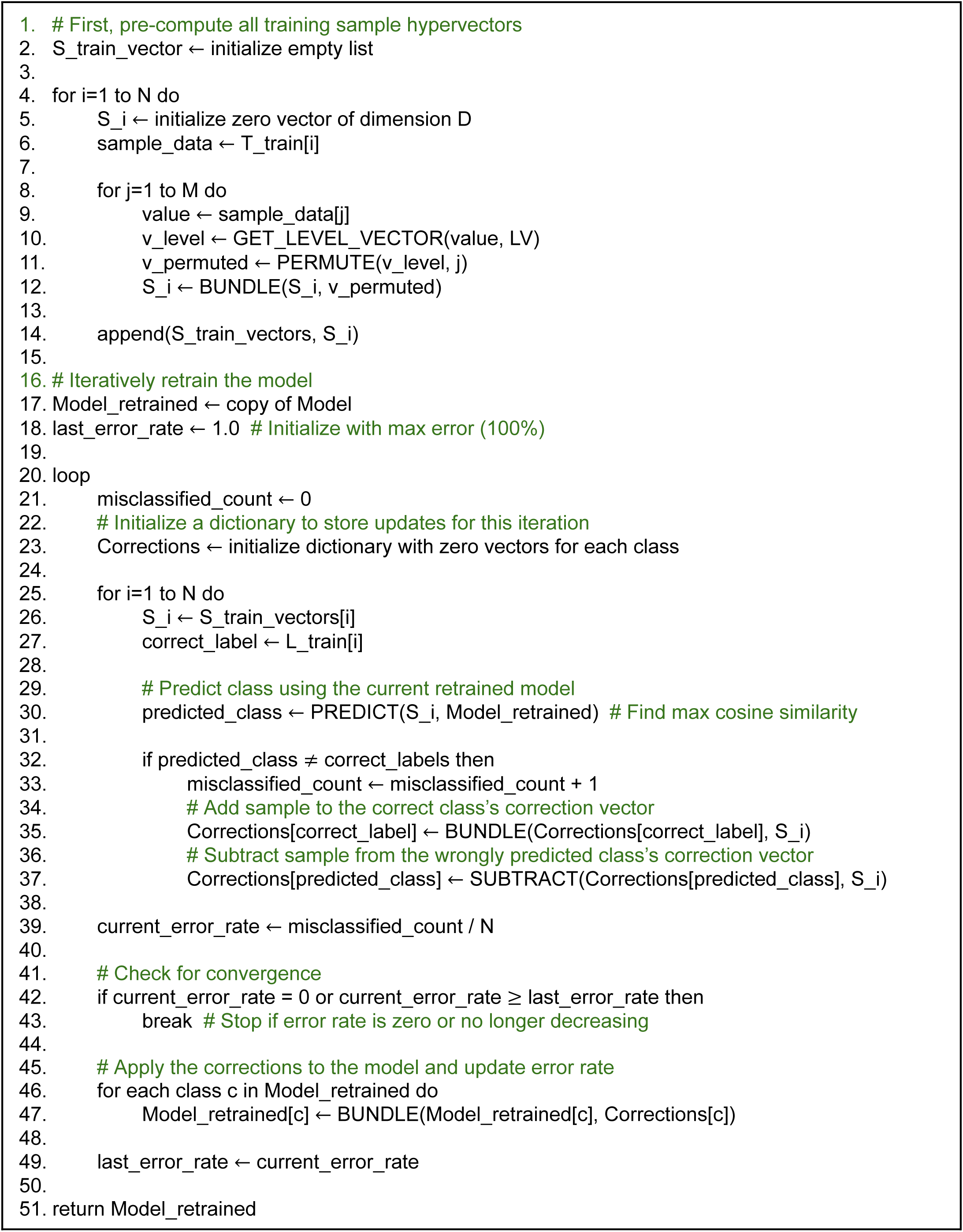

#### Stepwise regression

A critical challenge in applying machine learning to metagenomic data also lies in the identification of the most relevant features able to discriminate classes of samples and, eventually, for the investigation of potential biomarkers in the context of specific diseases and health conditions. While numerous feature selection methods exist for traditional machine learning models, to the best of our knowledge, no standardized techniques are currently available for classification models built according to the HDC paradigm. To address this gap, we introduced a stepwise regression approach as backward variable elimination, tailored specifically for feature selection in the context of HDC-based classification [25].

Backward variable elimination is an iterative feature selection technique [26] that starts producing *N* models with *N-1* features each, where *N* is the total number of features in the dataset. Each of these models are then evaluated according to a specific metric (e.g., accuracy, precision, recall, F1-score, etc.). Then, the process considers the models with the highest evaluation score only. It starts repeating iteratively the same process, considering, at every iteration, the set of features resulted from the intersection of the features used for building the best classification models only. The set of features gets smaller and smaller at every iteration until no features are left. Finally, an importance score can be assigned to each feature according to the iteration where it gets discarded. The last features to be discarded have a higher importance score.

## Results

Here, we examine the performances of our classification model (HDC) in comparison with four of the most popular classical supervised machine learning classification algorithms, i.e., Decision Tree (DTC), Logistic Regression (LRC), Random Forest (RFC), and Support Vector Machine (SVC). We also test our recursive backward variable elimination technique against the same kind of feature selection based on the aforementioned four classical machine learning methods. We rely on the F1-score as the evaluation metric for comparing the predictive power of all the classification models. The F1-score is often preferred over accuracy, especially when dealing with imbalanced datasets, because it provides a balanced measure of precision and recall. The analyses were performed in 5-fold cross-validation on a workstation equipped with Intel® Xeon® Platinum 8276L CPUs (2.20 GHz) and the NVIDIA A100 GPU.

Results are analyzed based on each of the metadata listed in the previous section. We summarize the results considering four different color-coded scenarios:

- Violet box – in case the HDC approach performs better than all the other classifiers;
- Red box – in case the HDC approach performs better than at least one of the other classifiers and performs comparably the same with respect to the remaining classifiers;
- Blue box – in case the HDC approach performs comparably the same with respect to the other classifiers;
- Black box – in case at least one classical classifier performed better than the HDC-based one.

We use a threshold of 0.05 to decide whether an algorithm performs better than another. If the values are within the threshold range, we consider the performance similar or comparable. However, we only mark a performance as better or worse if the difference between F1-scores is greater than the threshold. In summary, a threshold of 0.05 indicates that we only consider HDC to have higher performance if it shows a 5% improvement in its prediction accuracy, helping us in the identification of real instances where the HDC-base classifier performs better than the classical approaches.

Here, we report the results of the comparative analysis for each of the dataset stratified by the selected metadata, all resulting in binary classification tasks.

### Smoker

This category comprises a total of 475 samples from three different studies. In the *XieH_2016* [27] and *KeohaneDM_2020* [28] datasets, HDC outperformed all the other classical algorithms. HDC outperformed DTC, SVC, and RFC, and showed comparable performance to LRC for the *GhensiP_2019* [29] dataset (Figure 2).

**Figure 2.**
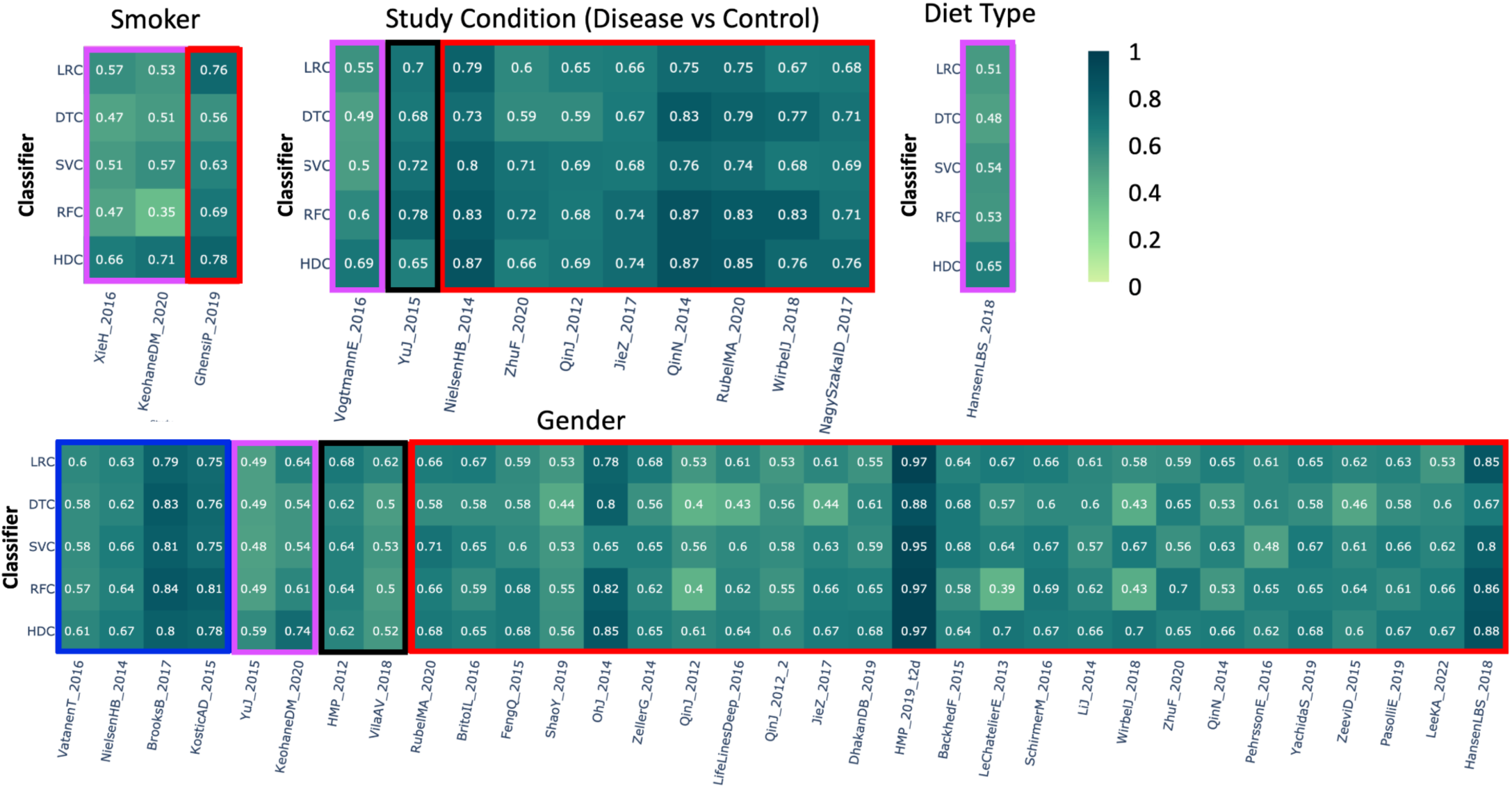
The heatmap represents the comparative performance of various classical and HDC classifiers applied to four different datasets: 1) Smoker vs. Non-smoker, 2) Study condition, 3) Diet type, and 4) Age group. The violet boxes indicate instances where the HDC classifier outperformed all classical algorithms. The red boxes indicate instances where the HDC classifier outperformed at least one classical algorithm. The blue boxes indicate instances where the HDC classifier performed comparably to the classical algorithms. The black boxes represent instances where at least one classifier outperformed the HDC classifier.

### Study conditions

This category includes datasets from 10 different studies, comprising 3,169 samples. Class labels represent various conditions versus control samples. In the heatmap, the first three columns show that the HDC performs comparably to the classical classifiers. In the fourth column, when applied to the *VogtmannE_2016* [30] dataset, HDC outperforms all other algorithms by a significant margin, represented with a violet box, achieving an F1 score of 0.69, while the other algorithms (LRC, DTC, SVC, and RFC) score 0.55, 0.49, 0.50, and 0.60, respectively. With the *YuJ_2015* [31] dataset, SVC and RFC classifiers perform better than HDC. However, the other two classical algorithms, LRC and DTC, show performances almost comparable to HDC. The red box around the results indicates that HDC outperformed at least one classical algorithm. Out of the 14 datasets, there are 19 instances where HDC outperforms at least one classical algorithm. In the first case, *NilsenHB_2014* [32], HDC showed significantly better performance than its classical counterparts, outperforming LRC, DTC, and SVC, while RFC showed comparable results. In the case of *ZhuF_2020* [33], RFC achieved a higher F1 score than HDC, while LRC and DTC showed significantly lower performance than HDC. SVC performed almost similarly to HDC. In the next case, *QinJ_2012* [34], HDC showed comparable performance to the other three classifiers, while it significantly outperformed LRC. In the case of *JieZ_2017* [35], HDC outperformed all the classical algorithms except RFC, with which it performed comparably, achieving an F1 score of 0.74. Similarly, with the *QinN_2014* dataset, HDC showed comparable performance to DTC and RFC, while outperforming LRC and SVC. In the case of *RubelMA_2020_A* [36], HDC performed comparably to RFC while significantly outperforming all the other classifiers. In the case of *WirbelJ_2018* [37], HDC showed comparable performance to DTC, outperformed LRC and SVC, while RFC outperformed HDC. Finally, in the case of *NagySzakaID_2017* [38], HDC performed comparably to DTC and RFC, while outperforming LRC and SVC classifiers (Figure 2).

### Diet type

In this category, the class labels represent high or low gluten-based diets. A total of 414 samples were included in this category, extracted from a single dataset, *HansenLBS_2018_B* [39]. Here, HDC outperformed all the other classifiers with an F1 score of 0.65 (Figure 2).

### Gender

In this category, the class labels represent two genders. This category includes around 17,692 samples from 33 different studies. In the case of *YuJ_2015* [31] and *KeohaneDM_2020* [28], the results show that HDC outperformed classical classifiers. However, in 4 out of 33 instances, HDC showed comparable performance to the other classifiers, while in 25 out of 33 instances, HDC outperformed at least one classical algorithm by a significant margin. The results also show that in 2 out of 33 instances, at least one classical algorithm outperformed HDC, while HDC showed comparable performance with some of the classical classifiers (Figure 2).

### Feeding practice

This category comprises a total of 151 samples from a single study, *YassourM_2018* [40]. In this dataset, the class labels represent either breastfeeding or mixed feeding practices. Results show that the HDC classifier performed better than DTC and SVC, while performing comparably to the other two classifiers, LRC and RFC (Figure 3).

**Figure 3.**
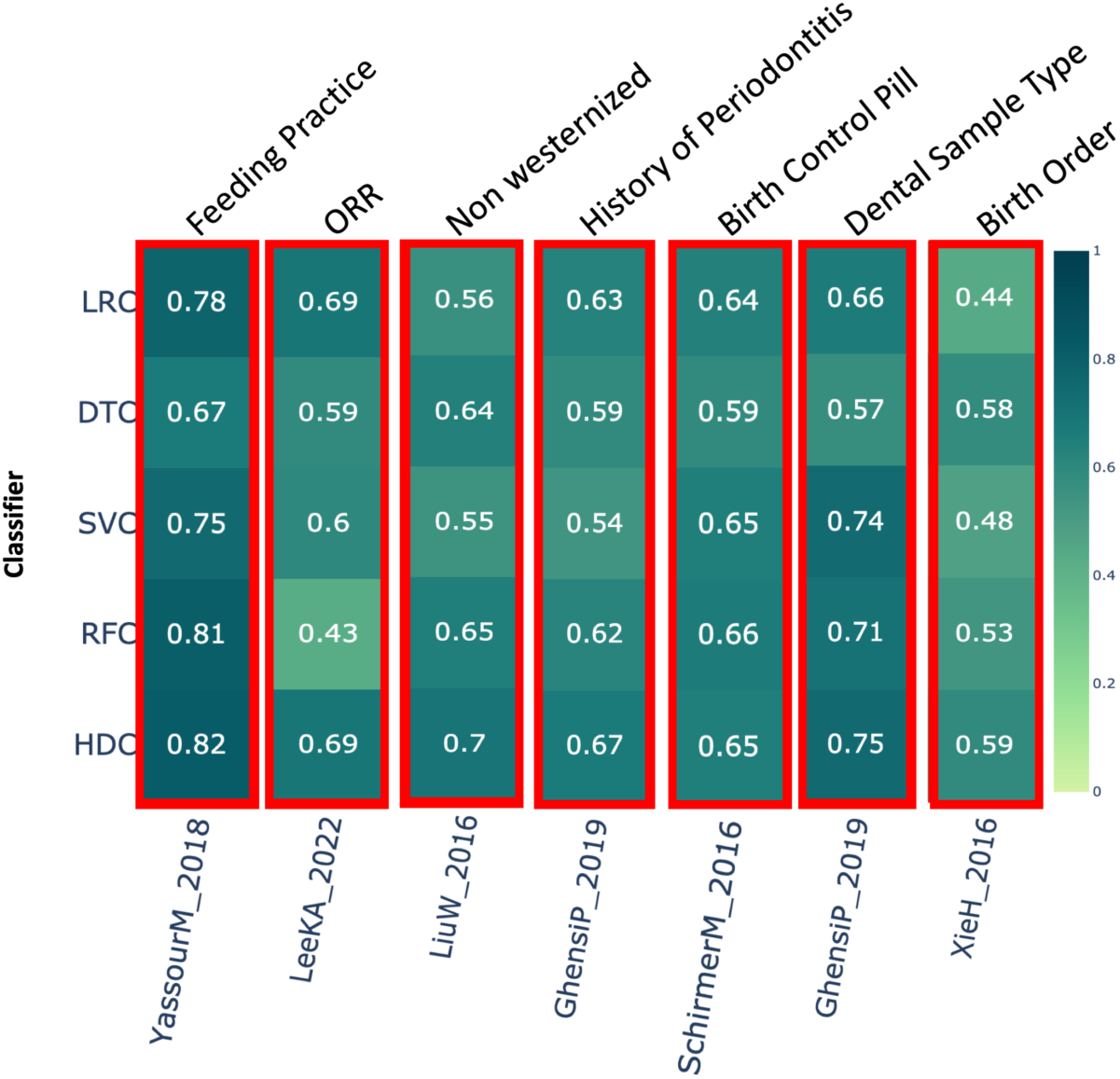
The heatmap shows the comparative performance of various classical and HDC classifiers applied to the following datasets: 1) Feeding practice, 2) ORR (Objective Response Rate), 3) Non-Westernized Lifestyle, 4) PSF12, 5) History of Periodontitis, 6) Birth Control Pill, 7) Dental Sample Type, and 8) Birth Order. The red boxes indicate instances where the HDC classifier outperformed at least one classical algorithm.

### ORR (Objective Response Rate)

The results show that HDC performed better than DTC, SVC, and RFC, while demonstrating comparable performance to LRC (Figure 3) on the dataset *LeeKA_2022* [41].

### Non-Westernized lifestyle

This category includes a total of 220 samples from two different studies. In the study *LiuW_2016* [42], HDC performed better than LRC and SVC while showing comparable performance to DTC and RFC (Figure 3).

### History of periodontitis

This category includes data from a single study, *GhensiP_2019* [29], which comprises a total 105 samples. Class label represents yes or no, indicating history of periodontitis in a patient. If HDC performs better then at least one classical classifier, represented with a red box around the results. The result shows that in this case HDC performs better than DTC and SVC and comparable to LRC and RFC (Figure 3).

### Birth control pill

This category includes data from a single study, *SchirmerM_2016* [43], and comprises around 286 samples. The class label represents whether birth control pills were used or not. In this case, HDC shows comparable performance with LRC, SVC, and RFC, while outperforming DTC (Figure 3).

### Dental sample type

This category includes 113 samples from a single dataset, *GhensiP_2019* [29]. The results suggest that HDC performed comparably to SVC and RFC, and outperformed LRC and DTC classifiers (Figure 3).

### Birth order

This category comprises a single study, *XieH_2016* [27], with 222 samples. Except for DTC, where HDC performed comparably, HDC outperformed all the other classical algorithms by a significant margin (Figure 3).

### Body site

This category includes around 312 samples from the dataset *BritoIL_2016* [44]. The class labels in this dataset represent various body sites for sample collection. In this case, the results show that all the algorithms performed similarly (Figure 4).

**Figure 4.**
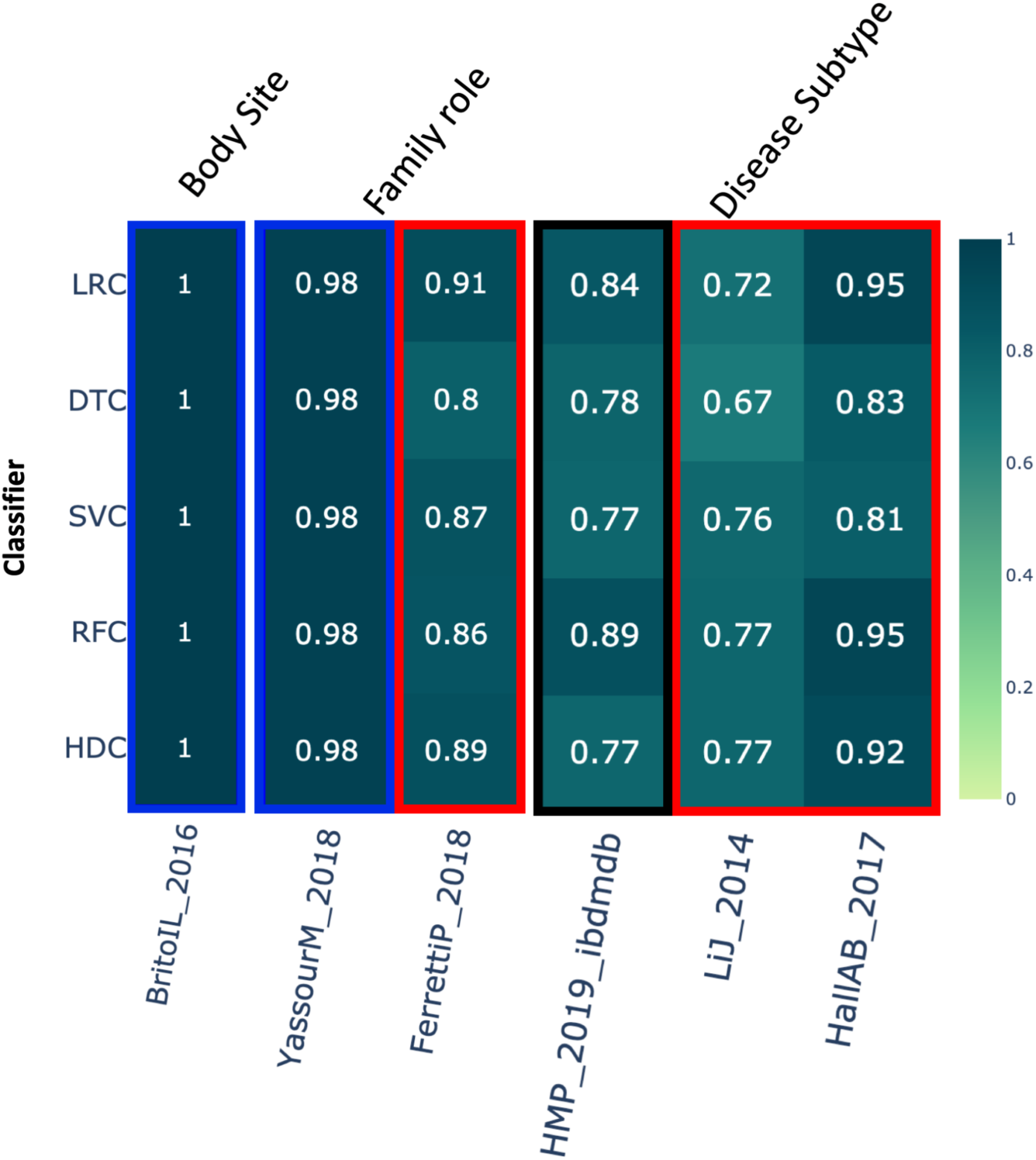
The heatmap shows the comparative performance of various classical and HDC classifiers applied to the following datasets: 1) Body site, 2) Family role, and 3) Disease subtype. The violet boxes indicate instances where the HDC classifier outperformed all classical algorithms. The red boxes indicate instances where the HDC classifier outperformed at least one classical algorithm. The blue boxes indicate instances where the HDC classifier performed comparably to the classical algorithms. The black boxes represent instances where at least one classifier outperformed the HDC classifier.

### Family role

This category includes a total of 485 samples based on two different studies. For the first dataset, *YassourM_2018* [40], HDC performed comparably to all the other algorithms. For the *FerrettiP_2018* dataset [45], HDC outperformed DTC while showing comparable performance to all the other algorithms (Figure 4).

### Disease subtype

This category includes data from 3 different studies with a total of 2,612 samples for classification. Class labels represent either Crohn’s disease or Ulcerative colitis. In these results, when we examine the first of the three datasets, *HMP_2019_ibdmdb*, RFC outperformed the HDC. However, HDC performed comparably to all the other classifiers. Conversely, in the case of the last two datasets, HDC performed better than at least one classifier. Specifically, in the *LiJ_2014* dataset [46], HDC outperformed the DTC, whereas in the *HallAB_2017* dataset [47], HDC outperformed both DTC and SVCs (Figure 4).

### Country

This category comprises a total number of 1,336 samples based on 3 different studies. In the first study, *PehrssonE_2016* [48], the result clearly shows that HDC performance is comparable to all the other algorithms. In the case of other three studies, in *CosteaPI_2017* [49] and *NielsenHB_2014* [32], as we can observe, HDC out performed DTC while performing comparable to other three classical classifiers (Figure 5).

**Figure 5.**
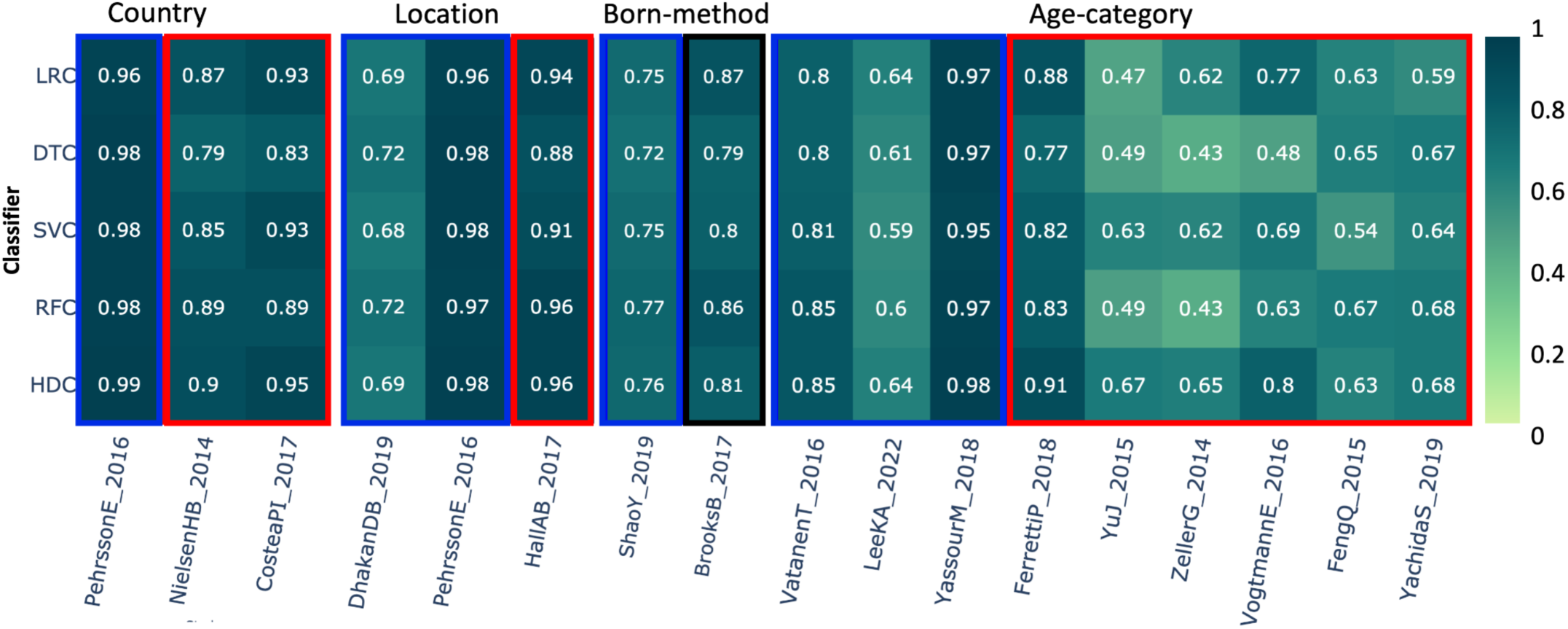
The heatmap shows the comparative performance of various classical and HDC classifiers applied to several datasets: 1) Country, 2) Location, 3) Born method, and 4) Age categories. The red boxes indicate instances where the HDC classifier outperformed at least one classical algorithm. The blue boxes indicate instances where the HDC classifier performed comparably to the classical algorithms. The black boxes represent instances where at least one classifier outperformed the HDC classifier.

### Location

This always appears in conjunction with the *Country of origin* category, and it is used to differentiate villages/towns/cities in the same country. This category includes a total of 1,010 samples from three different studies. In this dataset, the class labels represent various geographic locations around the world. For the first two datasets, *DhakanDB_2019* and *PehrssonE_2016*, HDC shows similar performance to the other four classical classifiers. In the *HallAB_2017* datasets [47], HDC outperformed SVC, but showed comparable performance to the other three classifiers (Figure 5).

### Born-method

This category includes data from two different studies and comprises more than 2,100 samples. In this dataset, class labels represent the birth method, which is either through a C-section or vaginal delivery. The blue box around the result shows that HDC classifiers perform comparably to all the classical algorithms, while the black box indicates that at least one classical classifier outperformed the HDC classifier. Results suggest that, in this category, the HDC classifier shows performance comparable to all the other classical classifiers when applied to the data derived from *ShaoY_2019* study [50]. However, when applied to the dataset from the other study *BrooksB_2017* [51], it shows performance similar to DTC, SVC, and RFC. However, LRC here performs better than the HDC classifier (Figure 5).

### Age-category

This category includes 3,137 samples from 9 different studies, with class labels representing different age groups. Results with the dataset show that HDC performed comparably to other classification algorithms for the *VatanenT_2016* [52], *LeekKA_2022* [41], and *YassourM_2018* [40] datasets. In the case of *FerrettiP_2018* [45], HDC outperformed DTC, SVC, and RFC while showing comparable performance to LRC. For the *YuJ_2015* dataset [31], HDC outperformed LRC, DTC, and RFC while performing comparably to SVC. HDC outperformed DTC and RFC for the *ZellerG_2014* dataset [53]. In the case of *VogtmannE_2016* [30], HDC outperformed DTC, SVC, and RFC while performing comparably to LRC. For the *FengQ_2015* dataset [54], HDC performed better than SVC while showing a comparable F1 score to other classifiers. Similarly, for the *YachidaS_2019* dataset [55], HDC showed comparable performance to DTC, SVC, and RFC while outperforming LRC (Figure 5).

Overall, there are various instances where HDC completely outperforms all the classical algorithms. For instance, in the cases of *gluten diet*, *study condition*, *smoker*, and *gender*, we observe 6 datasets where HDC outperforms all the classical algorithms. Predominantly, we also see instances (indicated with a red box) where HDC performs better than at least one classical algorithm while being comparable to others, demonstrating the superiority of HDC in several instances. In various categories the second predominant condition we can see where HDC shows equal performance, represented with a blue box, in comparison to classical classifiers. However, in various categories including *birth method*, *disease subtype*, *gender* and *study condition*, at least one of the classical algorithms outperforms the HDC classifier (black box).

We also examine the number of selected features based on various machine learning (ML) models. We find that all the algorithms show significant variability in the number of selected features. RFC and DTC have the highest means, 535 and 512 respectively, of selected features during different classification tasks. LRC also shows a comparatively high number of selected features with a mean of 303. On the other hand, SVC shows a lowest mean of 161 for the number of selected features across various classification tasks. HDC shows a mean of around 217 selected features across various classification tasks (Figure 6).

**Figure 6.**
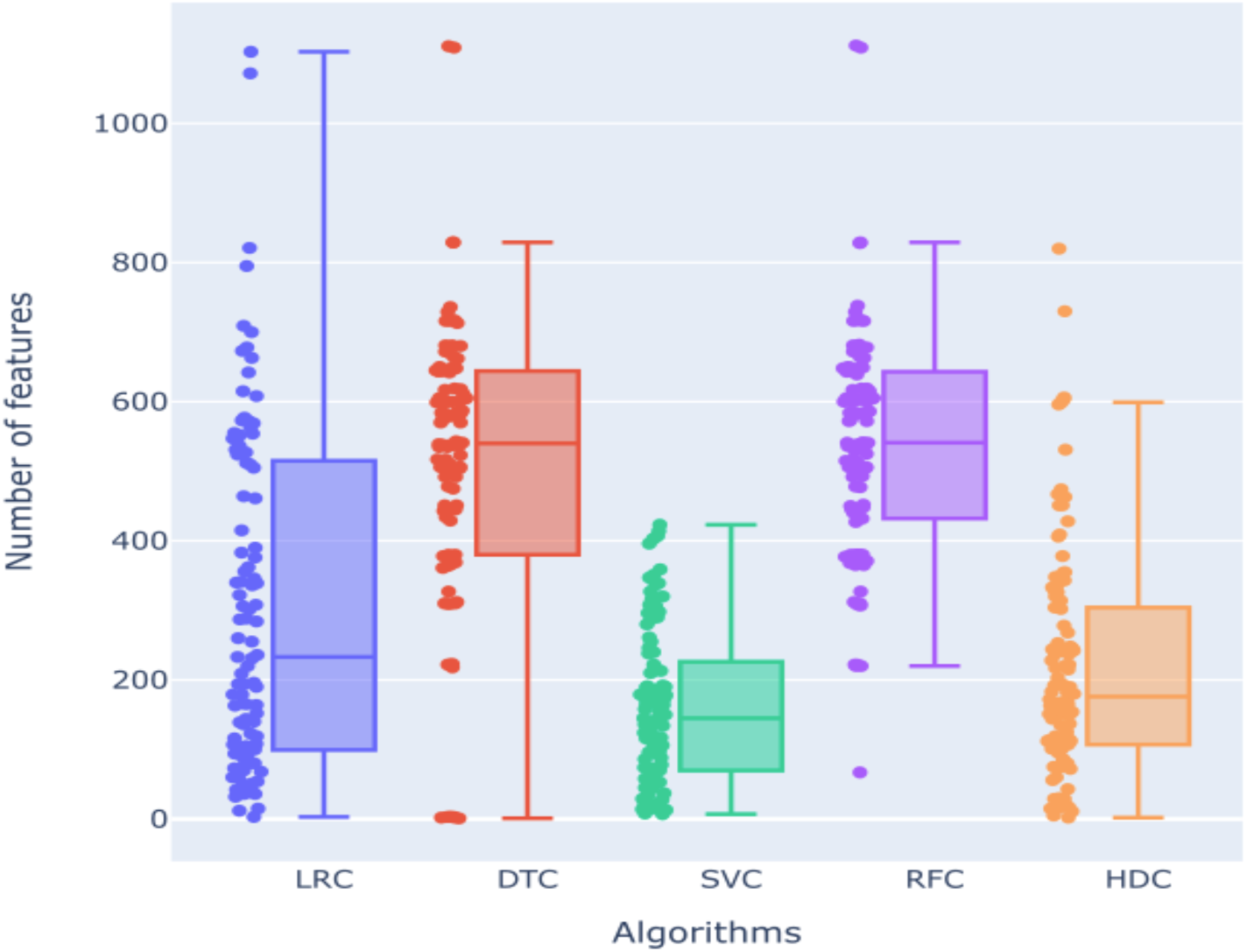
Distribution of the number of features selected by each of the algorithms (LRC, DTC, SVC, RFC, and HDC) considered in our comparative analysis.

Our analysis of feature selection across different classifiers exhibits distinct patterns in the number of features each method selects, which has implications for their performance and robustness. The Decision Tree Classifier (DTC) exhibited a wide range in the number of selected features, from as few as 1 to as many as 1,111 features, with an average of 512 features. This broad range suggests that DTC is highly variable in its feature selection, which may contribute to its inconsistency in performance. Similarly, the Logistic Regression Classifier (LRC) showed a range from 3 to 1,103 selected features, with a mean of 303. The lower mean compared to DTC indicates that LRC tends to select fewer features on average, which could be due to its reliance on linear combinations of features and regularization techniques that limit overfitting by reducing the number of selected features. The Random Forest Classifier (RFC) selected features within the range of 67 to 1,112, with a mean of 535. RFC’s higher minimum number of selected features and overall mean suggest that it tends to incorporate a substantial number of features to maintain its ensemble of decision trees, which might explain its robust performance across various datasets. In contrast, the Support Vector Classifier (SVC) demonstrated the narrowest range of selected features, with a minimum of 7 and a maximum of 423, and a mean of 161 features. This narrower range and lower mean indicate that SVC is more conservative in feature selection, likely due to its optimization for finding the best hyperplane that separates classes with the fewest features necessary. Finally, the HDC classifier selected features ranging from 2 to 820, with a mean of 217. HDC’s lower mean compared to other three algorithms and broader range compared to SVM suggest it is effective in identifying a smaller subset of relevant features, which may contribute to its competitive performance. The ability of HDC to perform well with fewer features indicates its potential for efficiency and reduced computational complexity. The higher performance of HDC on various datasets across various categories with minimal features highlights its potential as a versatile and efficient classifier. It maintains accuracy effectively even with a reduced feature set. Results demonstrate that while traditional classifiers like DTC, LRC, RFC, and SVC each have their strengths and weaknesses in feature selection, the HDC classifier stands out for its consistent performance with fewer selected features. Future work could explore optimizing feature selection methods further to enhance classifier performance and generalizability across different types of datasets.

Here, we focused on the cases where HDC outperformed all the classical algorithms and we further investigated the features selected by each method in these instances, Figure S6 (A-B). The mean number of features selected by HDC in these cases is 126.5, while for SVC, LRC, DTC, and RFC is 168.67, 249.5, 494.17, and 494.67 respectively, clearly highlighting the HDC ability in selecting smaller sets of relevant features maintaining high accuracy levels with respect to the classical counterpart. A paired t-test shows that the number of features selected by HDC is statistically significant according to a threshold of 0.05 on its p-value when compared to the number of features selected by DTC (*p*=1.6e-4) and RFC (*p*=1.6e-4). On the other hand, the comparison with SVC and LRC shows no statistical significance according to the same threshold, with a p-value of 2.318e-1 and 5.91e-2 respectively. However, the close-to-significance result between HDC and LRC suggests that HDC may still be more precise in selecting a relevant number of features compared to LRC. This reinforces the idea that HDC’s ability to select fewer features without sacrificing performance contributes to a better overall accuracy. Additionally, we also focused on the selected features in order to establish which method selected the best set of features able to significantly differentiate the two classes of samples in the context of those specific datasets. Here, we employed a paired Wilcoxon Rank-Sum test, finally computing the percentage of the statistically significant features according to the canonical threshold of 0.05 on the computed p-value. Our analysis revealed that HDC identified the highest number of relevant features (in percentage), with a mean of 6.68%, compared to the other methods, as shown in Figure S6 (C), clearly highlighting HDC’s ability of accurately selecting relevant features. Results of the Wilcoxon Rank-Sum test were reported in Spreadsheet S1.

Furthermore, among the features identified as statistically significant by the Wilcoxon Rank-Sum test, several well-established microbial biomarkers emerged in relation to the confounding variables used to stratify the metagenomic samples. For instance, *Eubacterium eligens*, *Coprococcus comes*, *Bifidobacterium adolescentis*, and *Anaerostipes hadrus*, all identified from the feature selection based on the HDC method on the *HansenLBS 2018* (Diet Type) dataset, have been previously associated with dietary habits [56–59]. Similarly, *Sellimonas intestinalis*, *Bifidobacterium pullorum*, and *Streptococcus australis* were selected from the datasets *XieH 2016* (Smoker) and *KeohaneDM 2020* (Smoker), belong to genera often used to discriminate between smoker and non-smoker individuals [60–63].

Finally, in terms of running time for the feature selection as backward variable elimination, HDC performed better than LRC and SVC, with a difference of 2,690 seconds and 18,784 seconds respectively on average (see Figure 7), approximately 45 minutes and 5.2 hours. On the other hand, it took 1,169 seconds and 954 seconds more on average compared to DTC and RFC, approximately 19.5 and 16 minutes respectively. However, these last two algorithms, as previously shown in Figure 6, selected a considerably larger set of features in general compared to HDC, which means that the backward variable elimination algorithm stopped working considerably earlier in these two cases, justifying their difference in running time compared to HDC. Spreadsheet S2 reports the detailed classification performances for each of the datasets here analyzed.

**Figure 7.**
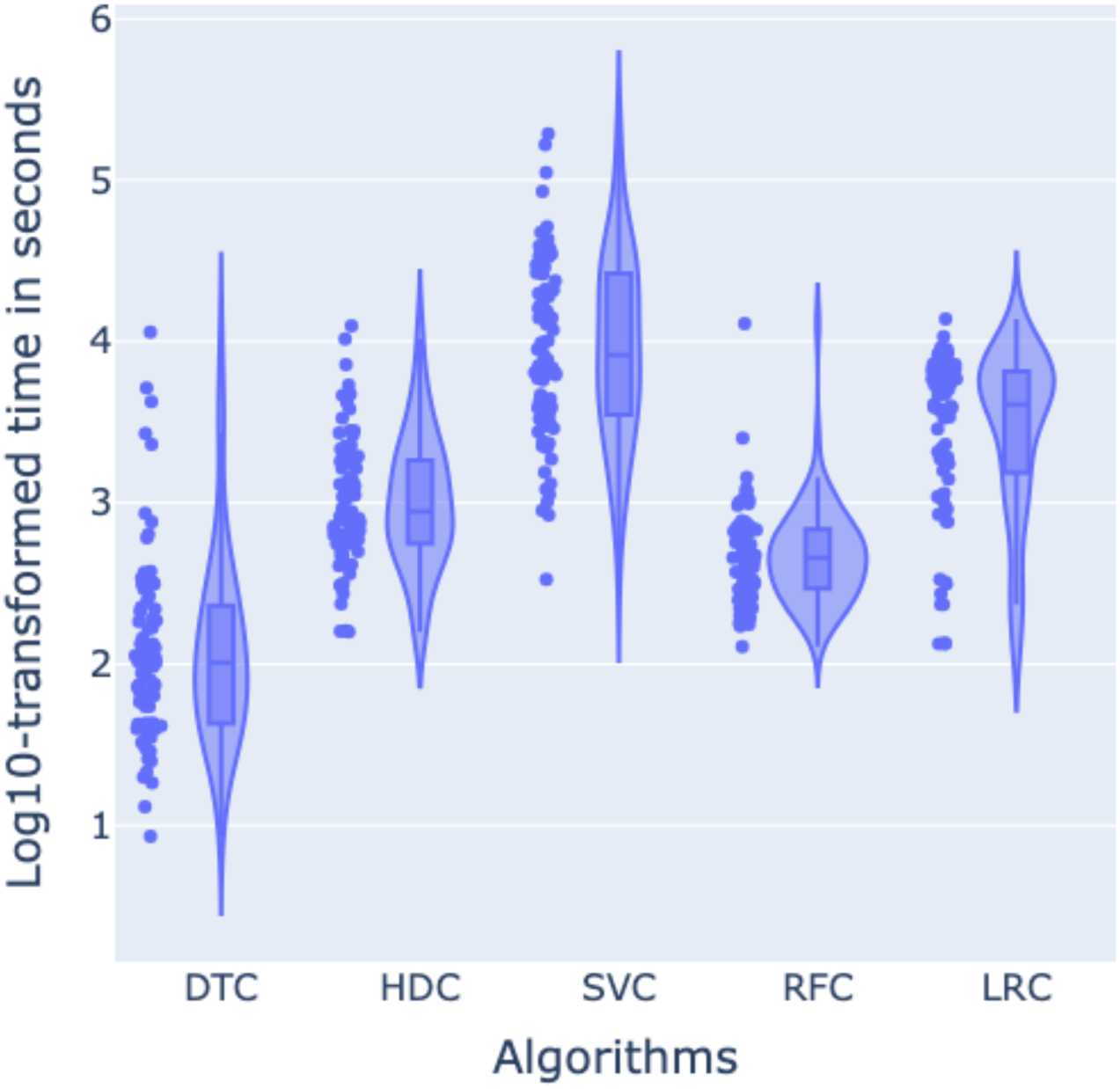
Distribution of log-transformed execution times (in seconds) for the feature selection tasks performed by five different machine learning models: DTC, HDC, SVC, RFC, and LRC. The violin plots display the kernel density estimation of log10-transformed times overlaid with individual data points, while the embedded boxplots indicate the median and interquartile range. This visualization highlights the variation in computation times across various ML models.

### Tools and Workflow

The entire analysis is supplied with the manuscript in the form of Galaxy tools and workflows. Galaxy is a browser-based platform that offers a user-friendly biomedical data analysis solution, which allows users to perform complex analyses using an intuitive GUI [64–67]. The toolset includes four tools, each described in detail in Table 2. The first tool in the workflow is MicrobiomeData, which is a data fetching tool, pulls the supplied data listed in Table 1. This tool offers various pre-computed data matrices such as microbial abundance and pathway abundance for the given datasets. The second tool in the workflow is the FeatureSelection tool, which provides robust options for selecting features based on supervised and unsupervised methods to build accurate models. The third tool, MLClassifiers, includes nine different machine learning algorithms for classification and offers various options for model evaluation, such as cross-validation and performance metrics like accuracy, precision, and F1 score. To analyze the results, we have included a plotting tool that generates various visualizations presented in this study.

**Table 2:**
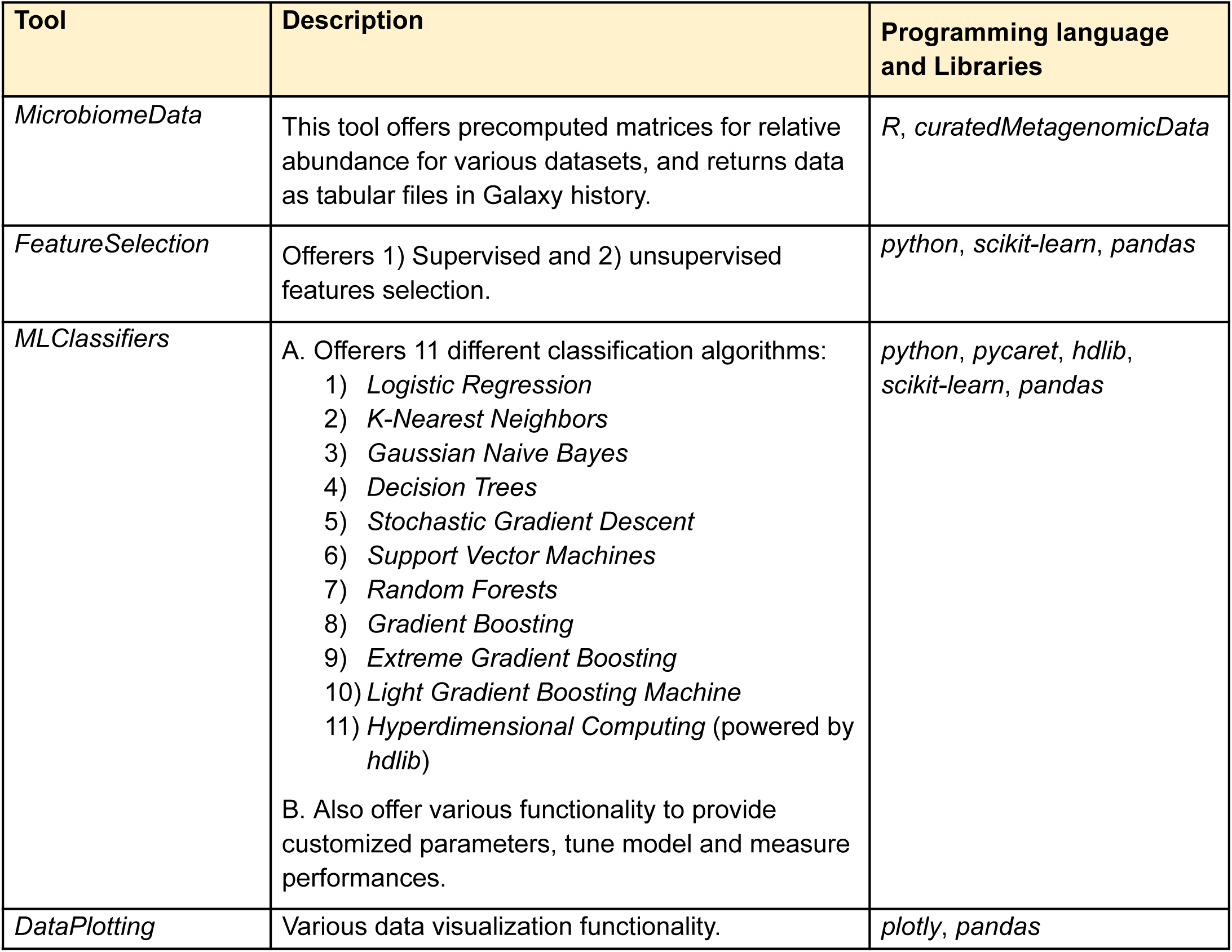
Steps of the Galaxy workflow with the involved tools and software libraries.

| Tool | Description | Programming language and Libraries |
| --- | --- | --- |
| *MicrobiomeData* | This tool offers precomputed matrices for relative abundance for various datasets, and returns data as tabular files in Galaxy history. | *R, curatedMetagenomicData* |
| *FeatureSelection* | Offers 1) Supervised and 2) unsupervised features selection. | *python, scikit-learn, pandas* |
| *MLClassifiers* | A. Offers 11 different classification algorithms:<br>1) *Logistic Regression*<br>2) *K-Nearest Neighbors*<br>3) *Gaussian Naive Bayes*<br>4) *Decision Trees*<br>5) *Stochastic Gradient Descent*<br>6) *Support Vector Machines*<br>7) *Random Forests*<br>8) *Gradient Boosting*<br>9) *Extreme Gradient Boosting*<br>10) *Light Gradient Boosting Machine*<br>11) *Hyperdimensional Computing* (powered by *hdlib*)<br>B. Also offer various functionality to provide customized parameters, tune model and measure performances. | *python, pycaret, hdlib, scikit-learn, pandas* |
| *DataPlotting* | Various data visualization functionality. | *plotly, pandas* |

Please note that, even though our tool provides an interface to 11 different classification algorithms (Hyperdimensional Computing (HDC), in addition to other 10 classical methods), our comparative analysis focused on the top performing ones, as mentioned in the Results section (LRC, DTC, SVC, and RFC).

## Discussion and Conclusions

These studies aimed to classify and understand the application of classical ML algorithms in the context of microbial abundance profiles. The primary purpose of this study is to evaluate the effectiveness of HDC classifiers in the classification of microbial abundance data and comparing HDC against most popular traditional classifiers including DTC, SVM, RFC and LRC. This comparative study aims to assess the efficacy, efficiency, and robustness of HDC classifiers against well-established and popular classical ML algorithms in handling the complex, high-dimensional nature of microbial abundance datasets, serving a crucial purpose in advancing understanding of microbiome analysis techniques. In this study, we seek to determine whether HDC classifiers can offer advantages such as improved classification accuracy, better handling of noise and variability inherent in microbiome data, faster processing times, or enhanced interpretability of results.

Our investigation into the application of a HDC-based supervised machine learning technique for classifying microbial profiles in metagenomic samples yielded promising results, demonstrating the potential of this novel computational paradigm to complement and, in some cases, surpass the performances of well established machine learning techniques like DT, LR, RF, and SVM.

The ability to efficiently handle and process this amount of data on a large-scale is due to its unique approach in representing information. HDC can indeed capture complex relationships within the data by leveraging high-dimensional vector and simple arithmetic operations. This efficiency makes this new kind of classification method an attractive alternative for analyzing large-scale metagenomic datasets for faster processing times and a more efficient use of computational resources.

Ultimately, this research contributes to the ongoing effort to refine and optimize tools for microbiome analysis, which is critical for advancing our understanding of microbial communities and their impact on various biological systems and environments.

When we compare the performance, HDC classifiers demonstrate consistent and competitive performance across a wide range of datasets, frequently outperforming or performing comparably to classical classifiers. In several instances, HDC outperformed all classical algorithms, as shown in the Results section. This is evident in various categories and conditions, such as Gluten Diet (1 out of 2), Study Condition (1 out of 14), Smoker (3 out of 5), and Gender (2 out of 44) marked with a violet box. On the other hand, as evident in various results, HDC outperformed some of the classical algorithms but performed comparable to others represented with red box. Specifically regarding gender, where HDC excelled despite being challenged by a large sample size (17692 samples belongs to 44 datasets), it consistently outperformed at least one classical classifier for 28 datasets while maintaining comparable performance with others. Similar results can be observed in the following categories: age (6 out of 10), birth control pill, birth order, country (3 out of 5), dental sample, disease subtype (2 out of 5), smoker (1 out of 4), family role (1 out of 2), feeding practice, history of periodontitis, location (2 out of 5), non-westernized lifestyle (2 out of 2), study condition (9 out of 14), and gluten-free diet (1 out of 2), where HDC outperformed at least one classical classifier while performing comparably to others. Interestingly, our findings underscored that the DTC classifier consistently exhibited the poorest performance among the classifiers evaluated, particularly in scenarios where HDC demonstrated clear superiority over classical methods. We also found that imbalanced data does not affect the overall performance of

HDC. In some cases, such as the *GhensiP_2019* and *XieH_2016* datasets in the smoker category, it even outperformed all classical classifiers. In other instances where the data was imbalanced, HDC still showed similar performance to other classifiers. This indicates that imbalanced data does not have a detrimental effect on the HDC classifier. This suggests that HDC may offer substantial advantages in accurately classifying and predicting outcomes across diverse datasets and categories compared to traditional classifiers. However, in various cases, HDC performed similarly or comparably to other classifiers, and we also observed instances where HDC was outperformed by classical classifiers.

There are a limited number of studies demonstrating the usability of ML algorithms for analyzing microbial abundance data, with existing ML techniques proving highly useful in this area [68,69]. However, in some cases classical algorithms often struggle with classification, explainability, and generalizability due to the complexity and experimental variability of microbiome datasets. This highlights the need for new and more advanced techniques. In this work, we present the application of a HDC classifier, showcasing its potential effectiveness in addressing challenges and underscoring the significance of this study in advancing ML applications for microbial analysis.

These results not only highlight HDC’s potential as a robust classifier in various research and clinical settings but also emphasize the importance of selecting appropriate ML models tailored to specific datasets and analytical goals. Further exploration and validation across larger and more diverse datasets would be beneficial to confirm and extend these findings, potentially paving the way for improved classification methodologies in genomic and clinical research.

It’s worth mentioning some of the limitations of this study. Microbial abundance data often comes from diverse sources, each with potential differences in quality, collection methods, and preprocessing. This variability can impact the robustness and generalizability of ML models, including the HDC one used in this study. While this study primarily focuses on comparing HDC to traditional methods, it lacks an in-depth comparison with more advanced approaches, such as deep learning methods, which have shown promise in analyzing complex microbial data despite their own limitations.

## Additional Information

### Availability

Microbial quantitative profiles pre-computed with *MetaPhlAn3* are all enclosed in the *curatedMetagenomicData* package for R available in Bioconductor at https://doi.org/doi:10.18129/B9.bioc.curatedMetagenomicData. The HDC-based supervised machine learning and feature selection technique is implemented in the *chopin2* [24] Python package and it is available in the Python Package Index (*pip install chopin2*) and Conda (*conda install -c conda-forge chopin2*). Its code is open-source and it is available on Github under the GPL-3.0 license at https://github.com/cumbof/chopin2. It makes use of the general purpose Python library for building vector-symbolic architectures *hdlib* [70] which is also available in the Python Package Index (*pip install hdlib*) and Conda (*conda install -c conda-forge hdlib*), while its code is open-source and available on GitHub under the MIT license at https://github.com/cumbof/hdlib. Classification results are available in Zenodo at https://doi.org/10.5281/zenodo.14872310. Galaxy [67,70] tools and workflow for reproducing the analysis results are available at https://github.com/jaidevjoshi83/MicroBiomML.

### Author Contributions

JJ and FC conceived the research, retrieved the datasets, defined the HDC classification models, and performed the comparative analysis; DB supervised the research and provided critical feedback for the definition of the manuscript; JJ, FC, and DB wrote the manuscript and agreed with its final version.

## Acknowledgments

We would like to thank Bryan Raubenolt from the Center for Computational Life Sciences of the Cleveland Clinic’s Lerner Research Institute for his constructive feedback that led to the final version of this manuscript.

We would also like to acknowledge the use of AI for language editing to improve clarity and readability, limited to sentence structure and word choice suggestions. Research ideas, data analysis, interpretations, and conclusions presented here are solely the product of authors’ own analysis and expertise.

## Supplementary Material

**Figure S1.**
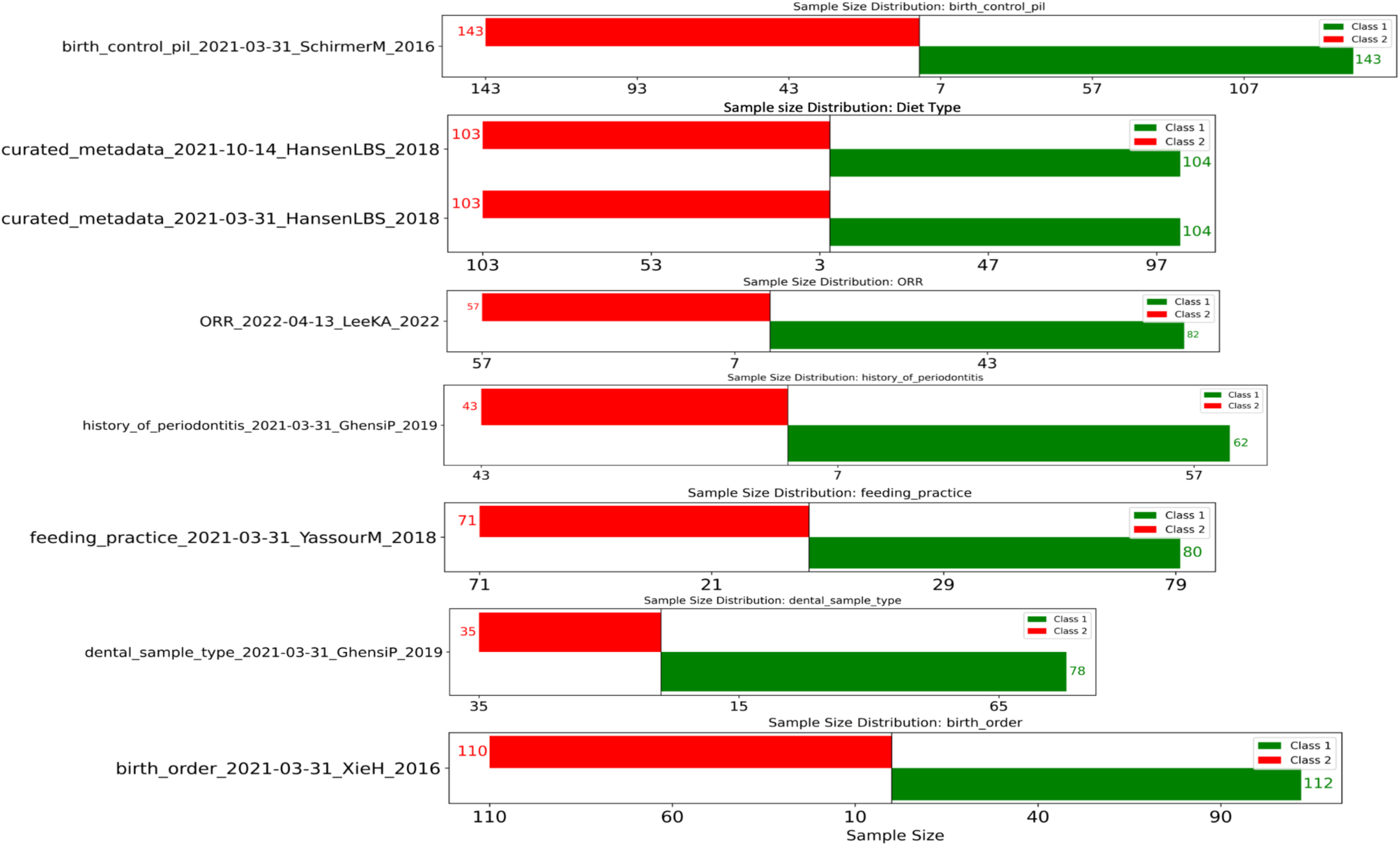
Sample distribution across the class labels: birth control pill vs. no birth control pill use.

**Figure S2.**
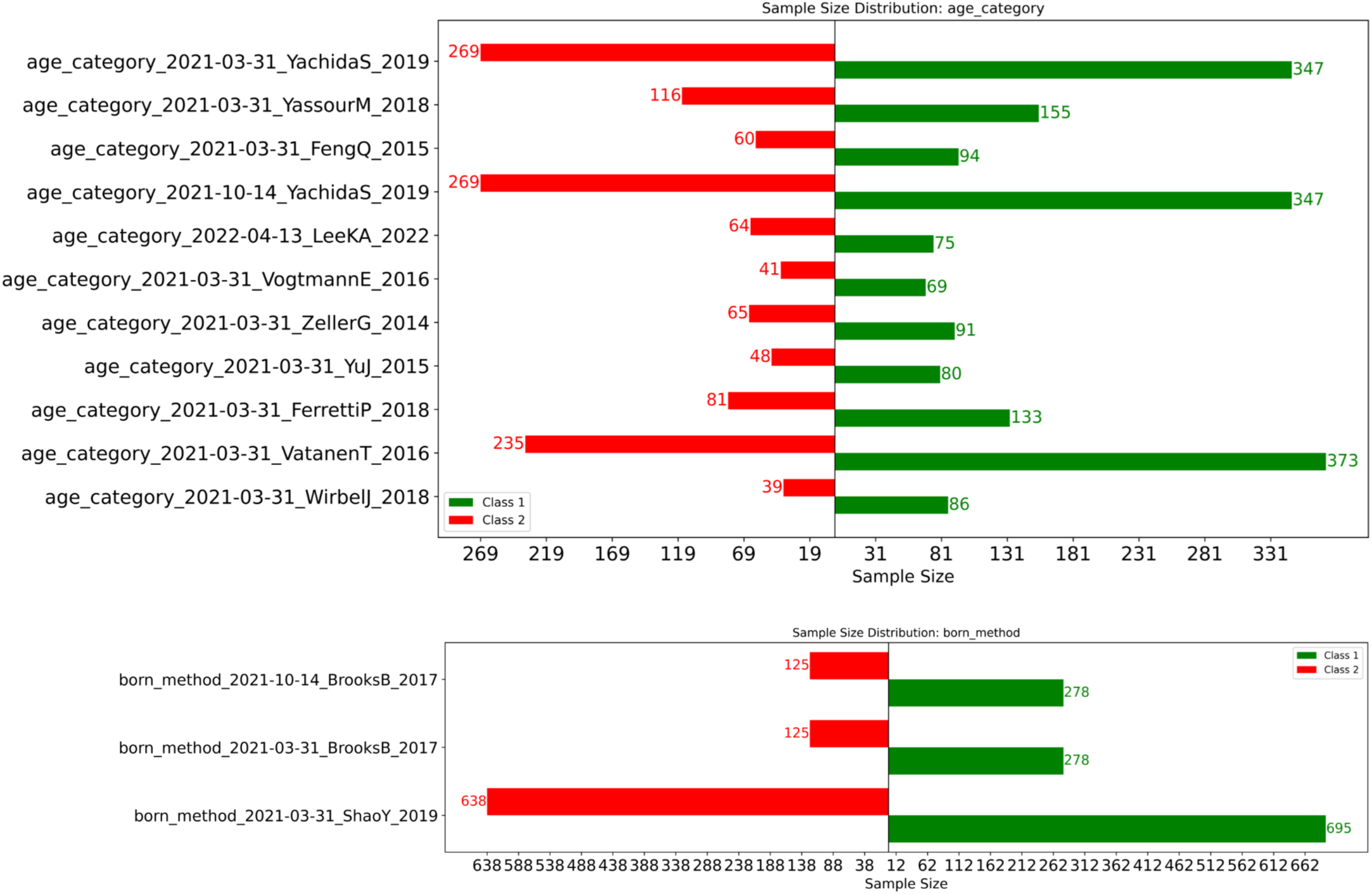
Sample distribution across both class labels in two different categories: age, where the class labels are young vs. old, and birth method, where the class labels are normal delivery vs. C-section.

**Figure S3.**
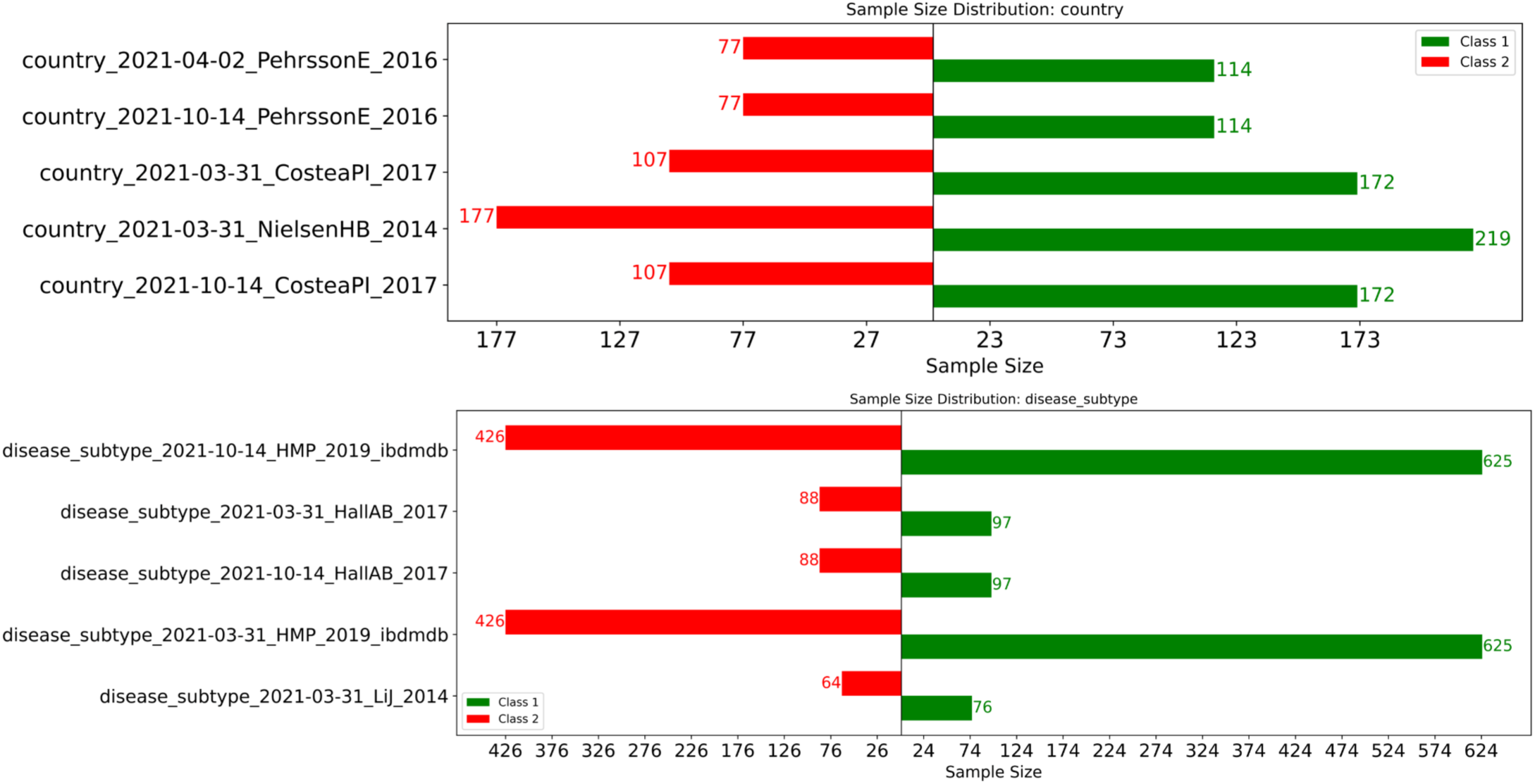
Sample distribution across two different categories: **country of origin**, where the class labels represent two different countries, and **disease subtype** where the class labels represent two different disease subtypes (e.g., Irritable Bowel Disease, Crohn’s Disease, or Diabetes Type I vs. Type II).

**Figure S4.**
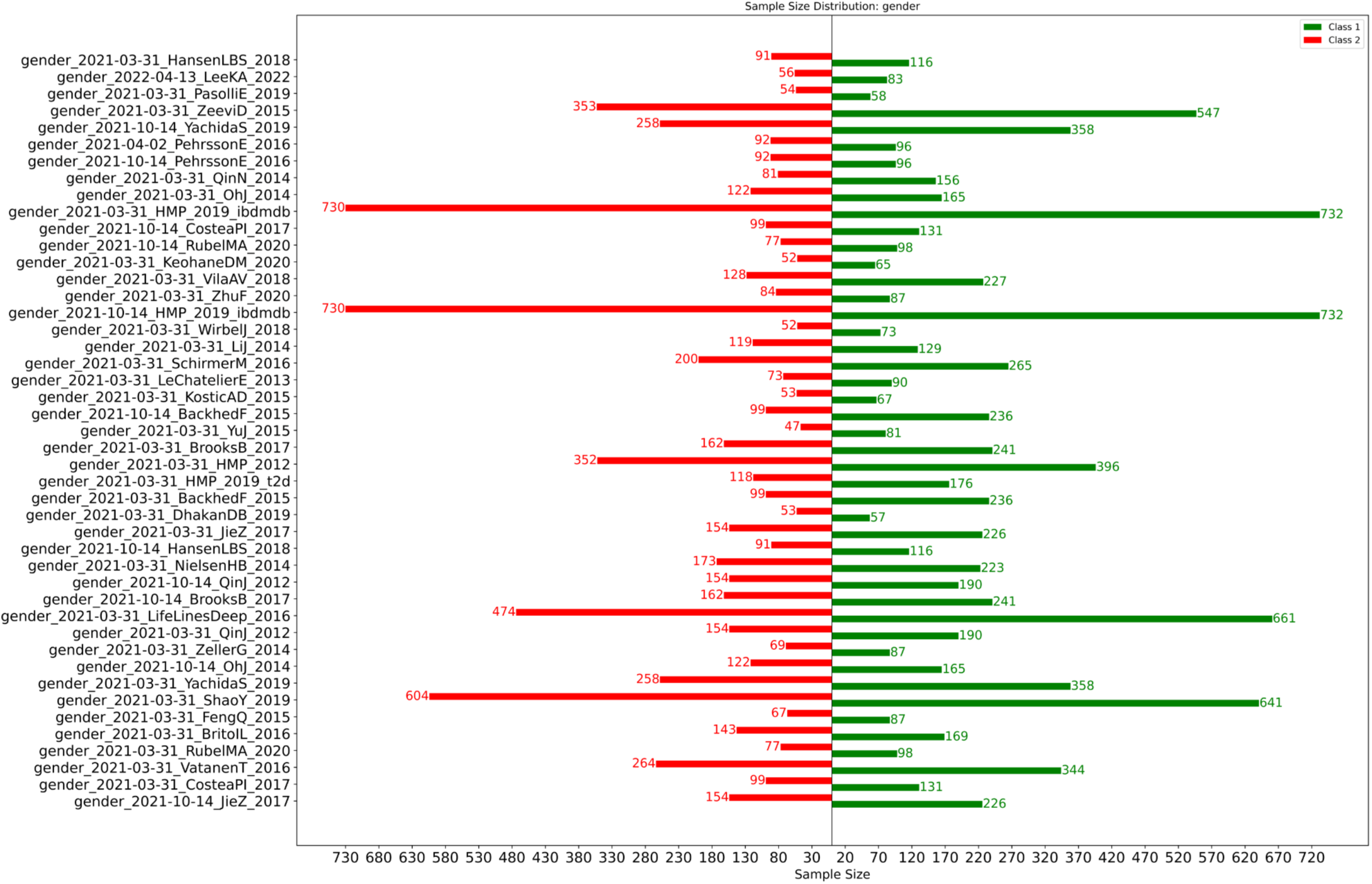
Sample distribution across both class labels male vs female.

**Figure S5.**
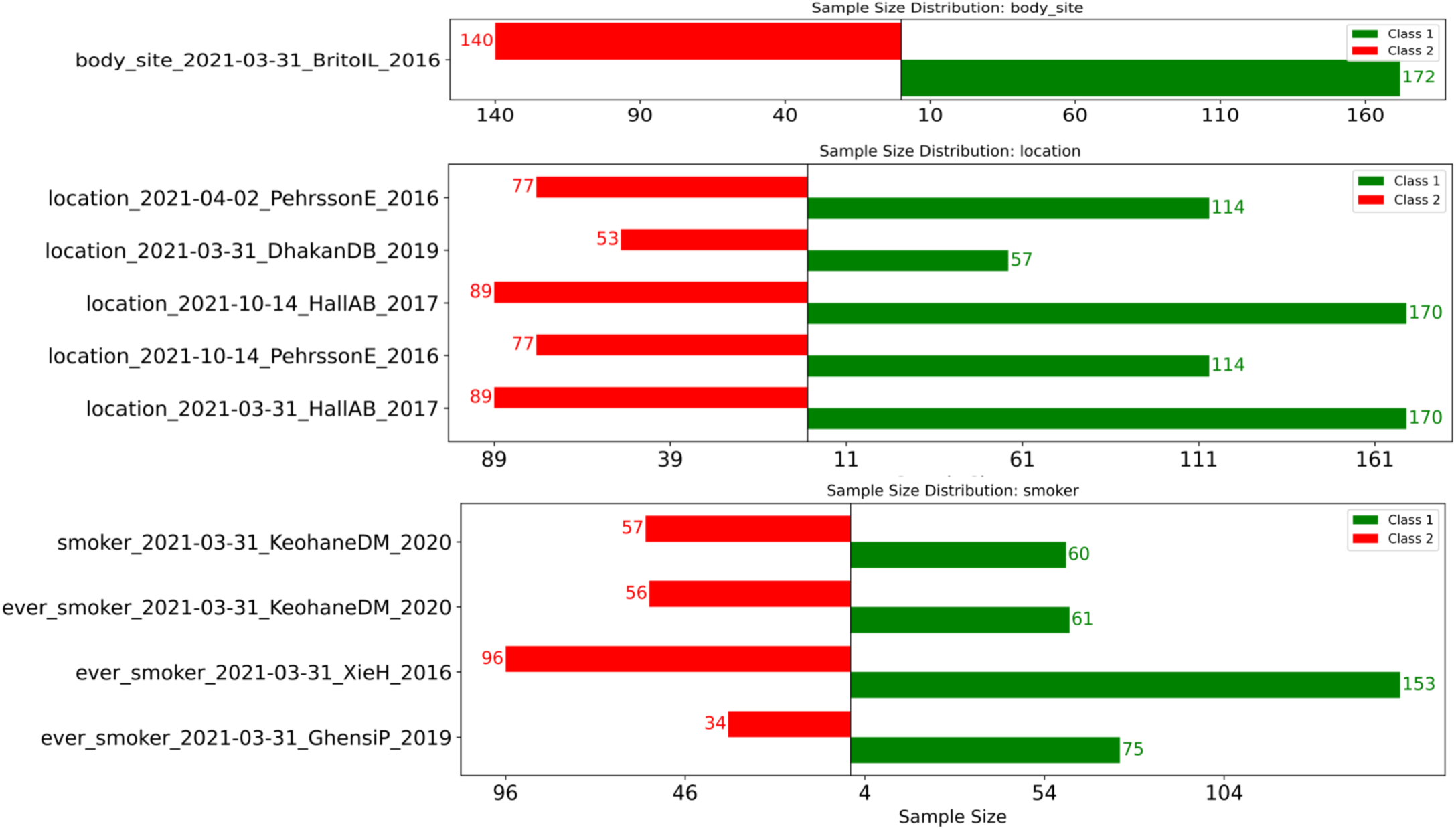
Sample distribution across three different categories: body site, where the class labels represent the different body parts where the samples were collected (e.g., skin, saliva, or feces); location, where the class labels correspond to two different cities; and smoking status, where the class labels are smoker vs. non-smoker.

**Figure S6.**
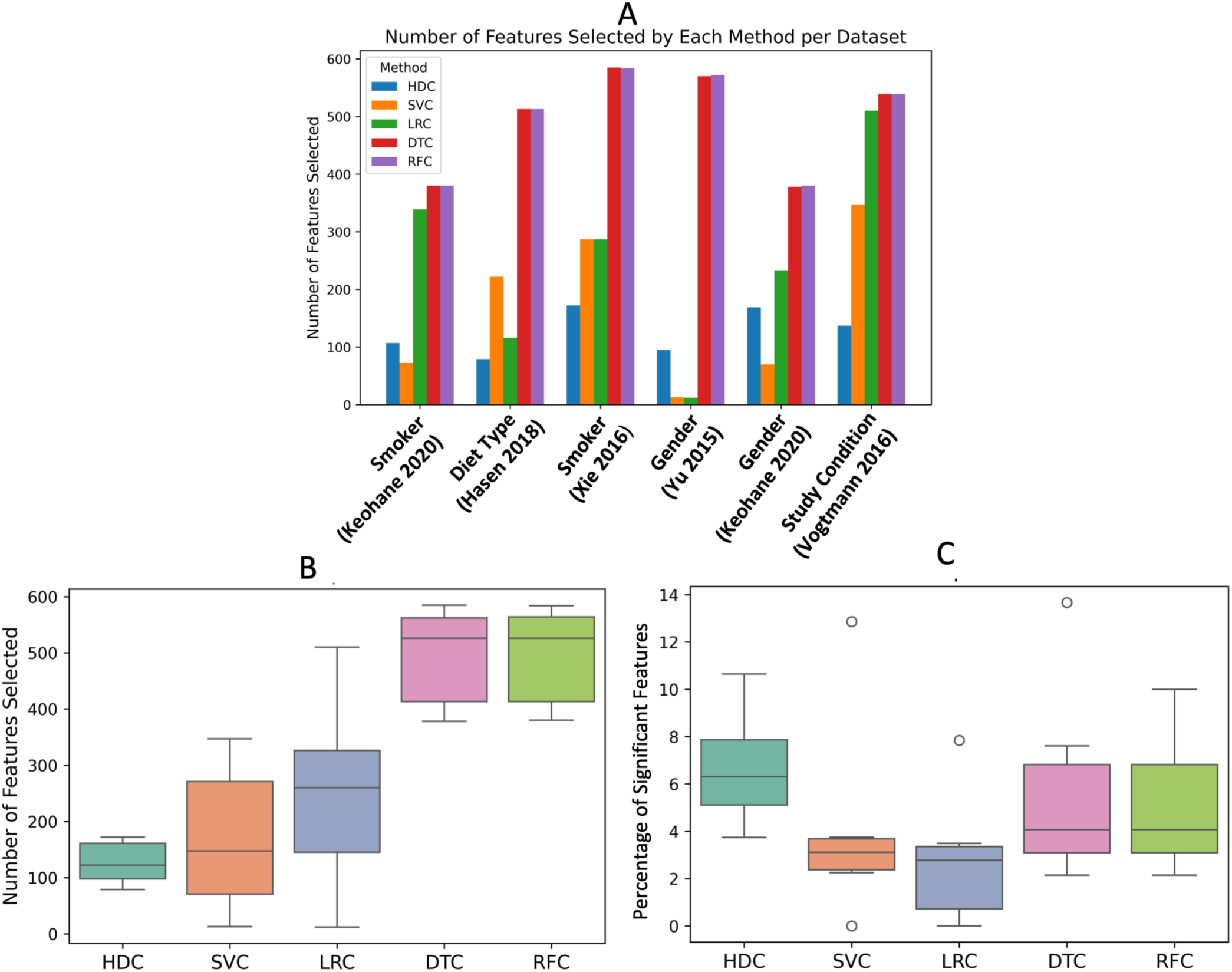
Panel A: The bar plot represents the number of selected features by each algorithm across the datasets for which HDC performs better in comparison to all the other algorithms. Panel B: The box plot shows the distribution of selected features across the same datasets considered in Panel A, where HDC outperformed all other algorithms. Panel C: Distribution of the percentage of significant features among selected features calculated based on the Wilcoxon Rank-Sum Test.

