## Supplementary material for "Large-scale classification of metagenomic samples: a comparative analysis of classical machine learning techniques vs a novel brain-inspired hyperdimensional computing approach": Spreadsheet S1

| HansenLBS 2018 – Diet Type | | | | | | | | | |
|---|---|---|---|---|---|---|---|---|---|
| **Hyperdimensional Computing** | | **Decision Trees** | | **Logistic Regression** | | **Random Forest** | | **Support Vector Machines** | |
| **Feature** | **p-value** | **Feature** | **p-value** | **Feature** | **p-value** | **Feature** | **p-value** | **Feature** | **p-value** |
| Eubacterium_eligens* | 0.0375 | Eubacterium_eligens | 0.0375 | Eubacterium_eligens | 0.0375 | Eubacterium_eligens | 0.0375 | Eubacterium_eligens | 0.0375 |
| Coprococcus_comes* | 0.0176 | Coprococcus_comes | 0.0176 | Coprococcus_comes | 0.0176 | Coprococcus_comes | 0.0176 | Coprococcus_comes | 0.0176 |
| Bifidobacterium_adolescentis* | 0.0036 | Bifidobacterium_adolescentis | 0.0036 | Bifidobacterium_adolescentis | 0.0036 | Bifidobacterium_adolescentis | 0.0036 | Bifidobacterium_adolescentis | 0.0036 |
| Anaerostipes_hadrus* | 0.0487 | Anaerostipes_hadrus | 0.0487 | | | Anaerostipes_hadrus | 0.0487 | Lactobacillus_gasseri | 0.0434 |
| | | Eisenbergiella_tayi | 0.0219 | | | Eisenbergiella_tayi | 0.0219 | Bifidobacterium_pseudolongum | 0.0455 |
| | | Streptococcus_sp_A12 | 0.0100 | | | Streptococcus_sp_A12 | 0.0100 | | |
| | | Lactobacillus_sanfranciscensis | 0.0000 | | | Lactobacillus_sanfranciscensis | 0.0000 | | |
| | | Aeriscardovia_aeriphila | 0.0136 | | | Aeriscardovia_aeriphila | 0.0136 | | |
| | | Clostridium_bolteae_CAG_59 | 0.0276 | | | Clostridium_bolteae_CAG_59 | 0.0276 | | |
| | | Lactobacillus_gasseri | 0.0434 | | | Lactobacillus_gasseri | 0.0434 | | |
| | | Bifidobacterium_pseudolongum | 0.0455 | | | Bifidobacterium_pseudolongum | 0.0455 | | |

| XieH 2016 – Smoker | | | | | | | | | |
|---|---|---|---|---|---|---|---|---|---|
| **Hyperdimensional Computing** | | **Decision Trees** | | **Logistic Regression** | | **Random Forest** | | **Support Vector Machines** | |
| **Feature** | **p-value** | **Feature** | **p-value** | **Feature** | **p-value** | **Feature** | **p-value** | **Feature** | **p-value** |
| Clostridium_leptum | 0.0148 | Bacteroides_clarus | 0.0165 | Methanobrevibacter_smithii | 0.0358 | Bacteroides_clarus | 0.0165 | Methanobrevibacter_smithii | 0.0358 |
| Catabacter_hongkongensis | 0.0205 | Agathobaculum_butyriciproducens | 0.0006 | Anaerotruncus_sp_CAG_528 | 0.0230 | Agathobaculum_butyriciproducens | 0.0006 | Anaerotruncus_sp_CAG_528 | 0.0230 |
| Methanobrevibacter_smithii | 0.0358 | Alistipes_shahii | 0.0134 | Bacteroides_sp_CAG_633 | 0.0224 | Alistipes_shahii | 0.0134 | Bacteroides_sp_CAG_633 | 0.0224 |
| Firmicutes_bacterium_CAG_238 | 0.0451 | Coprobacter_fastidiosus | 0.0269 | Firmicutes_bacterium_CAG_424 | 0.0370 | Coprobacter_fastidiosus | 0.0269 | Firmicutes_bacterium_CAG_424 | 0.0370 |
| Mogibacterium_timidum* | 0.0285 | Clostridium_leptum | 0.0148 | Sellimonas_intestinalis | 0.0237 | Clostridium_leptum | 0.0148 | Sellimonas_intestinalis | 0.0237 |
| Actinomyces_viscosus* | 0.0342 | Enorma_massiliensis | 0.0373 | Lactobacillus_salivarius | 0.0007 | Enorma_massiliensis | 0.0373 | Lactobacillus_salivarius | 0.0007 |
| Clostridium_saccharolyticum | 0.0165 | Catabacter_hongkongensis | 0.0205 | Anaerofustis_stercorihominis | 0.0415 | Catabacter_hongkongensis | 0.0205 | Anaerofustis_stercorihominis | 0.0415 |
| Sellimonas_intestinalis* | 0.0237 | Dorea_sp_CAG_317 | 0.0243 | Lactobacillus_brevis | 0.0285 | Dorea_sp_CAG_317 | 0.0243 | Lactobacillus_brevis | 0.0285 |
| Bifidobacterium_pullorum* | 0.0285 | Methanobrevibacter_smithii | 0.0358 | Bifidobacterium_pullorum | 0.0285 | Methanobrevibacter_smithii | 0.0358 | Bifidobacterium_pullorum | 0.0285 |
| | | Firmicutes_bacterium_CAG_238 | 0.0451 | Enterococcus_casseliflavus | 0.0285 | Firmicutes_bacterium_CAG_238 | 0.0451 | Enterococcus_casseliflavus | 0.0285 |
| | | Blautia_sp_CAG_257 | 0.0400 | | | Blautia_sp_CAG_257 | 0.0400 | | |
| | | Veillonella_rogosae | 0.0118 | | | Veillonella_rogosae | 0.0118 | | |
| | | Anaerotruncus_sp_CAG_528 | 0.0230 | | | Anaerotruncus_sp_CAG_528 | 0.0230 | | |
| | | Fusobacterium_sp_CAG_439 | 0.0090 | | | Fusobacterium_sp_CAG_439 | 0.0090 | | |
| | | Dorea_sp_D27 | 0.0342 | | | Dorea_sp_D27 | 0.0342 | | |
| | | Mogibacterium_timidum | 0.0285 | | | Mogibacterium_timidum | 0.0285 | | |
| | | Actinomyces_viscosus | 0.0342 | | | Actinomyces_viscosus | 0.0342 | | |
| | | Bacteroides_sp_CAG_633 | 0.0224 | | | Bacteroides_sp_CAG_633 | 0.0224 | | |
| | | Clostridium_saccharolyticum | 0.0165 | | | Clostridium_saccharolyticum | 0.0165 | | |
| | | Firmicutes_bacterium_CAG_424 | 0.0370 | | | Firmicutes_bacterium_CAG_424 | 0.0370 | | |
| | | Sellimonas_intestinalis | 0.0237 | | | Sellimonas_intestinalis | 0.0237 | | |
| | | Lactobacillus_salivarius | 0.0007 | | | Lactobacillus_salivarius | 0.0007 | | |
| | | Anaerofustis_stercorihominis | 0.0415 | | | Anaerofustis_stercorihominis | 0.0415 | | |
| | | Lactobacillus_brevis | 0.0285 | | | Lactobacillus_brevis | 0.0285 | | |
| | | Bifidobacterium_pullorum | 0.0285 | | | Bifidobacterium_pullorum | 0.0285 | | |
| | | Enterococcus_casseliflavus | 0.0285 | | | Enterococcus_casseliflavus | 0.0285 | | |

| YuJ 2015 – Gender | | | | | | | | | |
|---|---|---|---|---|---|---|---|---|---|
| **Hyperdimensional Computing** | | **Decision Trees** | | **Logistic Regression** | | **Random Forest** | | **Support Vector Machines** | |
| **Feature** | **p-value** | **Feature** | **p-value** | **Feature** | **p-value** | **Feature** | **p-value** | **Feature** | **p-value** |
| Gordonibacter_pamelaeae | 0.0111 | Firmicutes_bacterium_CAG_83 | 0.0450 | | | Firmicutes_bacterium_CAG_83 | 0.0450 | | |
| Dielma_fastidiosa | 0.0202 | Gordonibacter_pamelaeae | 0.0111 | | | Gordonibacter_pamelaeae | 0.0111 | | |
| Ruminococcaceae_bacterium_D16 | 0.0146 | Actinomyces_sp_ICM47 | 0.0187 | | | Actinomyces_sp_ICM47 | 0.0187 | | |
| Clostridium_lavalense | 0.0343 | Eisenbergiella_tayi | 0.0063 | | | Eisenbergiella_tayi | 0.0063 | | |
| Actinomyces_graevenitzii | 0.0273 | Eisenbergiella_massiliensis | 0.0385 | | | Eisenbergiella_massiliensis | 0.0385 | | |
| Streptococcus_mitis | 0.0213 | Dielma_fastidiosa | 0.0202 | | | Dielma_fastidiosa | 0.0202 | | |
| Streptococcus_downei | 0.0452 | Clostridium_bolteae | 0.0227 | | | Clostridium_bolteae | 0.0227 | | |
| | | Ruminococcaceae_bacterium_D16 | 0.0146 | | | Ruminococcaceae_bacterium_D16 | 0.0146 | | |
| | | Streptococcus_infantis | 0.0009 | | | Streptococcus_infantis | 0.0009 | | |
| | | Clostridium_lavalense | 0.0343 | | | Clostridium_lavalense | 0.0343 | | |
| | | Actinomyces_graevenitzii | 0.0273 | | | Actinomyces_graevenitzii | 0.0273 | | |
| | | Roseburia_sp_CAG_309 | 0.0175 | | | Roseburia_sp_CAG_309 | 0.0175 | | |
| | | Streptococcus_mitis | 0.0213 | | | Streptococcus_mitis | 0.0213 | | |
| | | Enterococcus_faecalis | 0.0311 | | | Enterococcus_faecalis | 0.0311 | | |
| | | Abiotrophia_sp_HMSC24B09 | 0.0396 | | | Abiotrophia_sp_HMSC24B09 | 0.0396 | | |
| | | Lactobacillus_paragasseri | 0.0017 | | | Lactobacillus_paragasseri | 0.0017 | | |
| | | Fusobacterium_ulcerans | 0.0032 | | | Fusobacterium_ulcerans | 0.0032 | | |
| | | Bacteroides_sp_OM08_11 | 0.0059 | | | Bacteroides_sp_OM08_11 | 0.0059 | | |
| | | Oribacterium_parvum | 0.0396 | | | Oribacterium_parvum | 0.0396 | | |
| | | Oribacterium_sinus | 0.0127 | | | Oribacterium_sinus | 0.0127 | | |
| | | Streptococcus_downei | 0.0452 | | | Streptococcus_downei | 0.0452 | | |

| VogtmannE 2016 – Study Condition | | | | | | | | | |
|---|---|---|---|---|---|---|---|---|---|
| **Hyperdimensional Computing** | | **Decision Trees** | | **Logistic Regression** | | **Random Forest** | | **Support Vector Machines** | |
| **Feature** | **p-value** | **Feature** | **p-value** | **Feature** | **p-value** | **Feature** | **p-value** | **Feature** | **p-value** |
| Firmicutes_bacterium_CAG_95 | 0.0394 | Ruminococcus_gnavus | 0.0183 | Ruminococcus_gnavus | 0.0183 | Ruminococcus_gnavus | 0.0183 | Firmicutes_bacterium_CAG_110 | 0.0035 |
| Firmicutes_bacterium_CAG_110 | 0.0035 | Roseburia_hominis | 0.0418 | Roseburia_hominis | 0.0418 | Roseburia_hominis | 0.0418 | Bacteroides_fragilis | 0.0354 |
| Escherichia_coli | 0.0028 | Eisenbergiella_tayi | 0.0314 | Eisenbergiella_tayi | 0.0314 | Eisenbergiella_tayi | 0.0314 | Porphyromonas_asaccharolytica | 0.0028 |
| Clostridium_symbiosum | 0.0049 | Firmicutes_bacterium_CAG_95 | 0.0394 | Firmicutes_bacterium_CAG_95 | 0.0394 | Firmicutes_bacterium_CAG_95 | 0.0394 | Clostridium_scindens | 0.0074 |
| Bacteroides_fragilis | 0.0354 | Firmicutes_bacterium_CAG_110 | 0.0035 | Firmicutes_bacterium_CAG_110 | 0.0035 | Firmicutes_bacterium_CAG_110 | 0.0035 | Parvimonas_micra | 0.0196 |
| Fusobacterium_nucleatum | 0.0003 | Asaccharobacter_celatus | 0.0363 | Asaccharobacter_celatus | 0.0363 | Asaccharobacter_celatus | 0.0363 | Dialister_pneumosintes | 0.0010 |
| Peptostreptococcus_stomatis | 0.0076 | Escherichia_coli | 0.0028 | Escherichia_coli | 0.0028 | Escherichia_coli | 0.0028 | Firmicutes_bacterium_CAG_534 | 0.0434 |
| Clostridium_scindens | 0.0074 | Flavonifractor_sp_An100 | 0.0271 | Flavonifractor_sp_An100 | 0.0271 | Flavonifractor_sp_An100 | 0.0271 | Bacteroides_pectinophilus | 0.0185 |
| Dialister_pneumosintes | 0.0010 | Clostridium_symbiosum | 0.0049 | Clostridium_symbiosum | 0.0049 | Clostridium_symbiosum | 0.0049 | Streptococcus_gallolyticus | 0.0124 |
| Bacteroides_pectinophilus | 0.0185 | Bacteroides_fragilis | 0.0354 | Bacteroides_fragilis | 0.0354 | Bacteroides_fragilis | 0.0354 | Methanosphaera_stadtmanae | 0.0231 |
| Streptococcus_gallolyticus | 0.0124 | Actinomyces_sp_HPA0247 | 0.0249 | Actinomyces_sp_HPA0247 | 0.0249 | Actinomyces_sp_HPA0247 | 0.0249 | Megamonas_funiformis_CAG_377 | 0.0434 |
| | | Oscillibacter_sp_PC13 | 0.0243 | Oscillibacter_sp_PC13 | 0.0243 | Oscillibacter_sp_PC13 | 0.0243 | Peptostreptococcus_anaerobius | 0.0124 |
| | | Clostridium_sp_CAG_167 | 0.0476 | Clostridium_sp_CAG_167 | 0.0476 | Clostridium_sp_CAG_167 | 0.0476 | Eubacterium_infirmum | 0.0434 |
| | | Roseburia_sp_CAG_309 | 0.0115 | Roseburia_sp_CAG_309 | 0.0115 | Gemella_haemolysans | 0.0326 | | |
| | | Gemella_haemolysans | 0.0326 | Gemella_haemolysans | 0.0326 | Fretibacterium_fastidiosum | 0.0361 | | |
| | | Fretibacterium_fastidiosum | 0.0361 | Fretibacterium_fastidiosum | 0.0361 | Mogibacterium_diversum | 0.0334 | | |
| | | Mogibacterium_diversum | 0.0334 | Mogibacterium_diversum | 0.0334 | Christensenella_minuta | 0.0352 | | |
| | | Christensenella_minuta | 0.0352 | Christensenella_minuta | 0.0352 | Bacteroides_intestinalis | 0.0080 | | |
| | | Bacteroides_intestinalis | 0.0080 | Bacteroides_intestinalis | 0.0080 | Prevotella_intermedia | 0.0434 | | |
| | | Prevotella_intermedia | 0.0434 | Prevotella_intermedia | 0.0434 | Gemella_morbillorum | 0.0115 | | |
| | | Gemella_morbillorum | 0.0115 | Solobacterium_moorei | 0.0014 | Solobacterium_moorei | 0.0014 | | |
| | | Solobacterium_moorei | 0.0014 | Anaerococcus_vaginalis | 0.0035 | Anaerococcus_vaginalis | 0.0035 | | |
| | | Anaerococcus_vaginalis | 0.0035 | Parvimonas_sp_KA00067 | 0.0231 | Parvimonas_sp_KA00067 | 0.0231 | | |
| | | Parvimonas_sp_KA00067 | 0.0231 | Peptoniphilus_lacrimalis | 0.0434 | Peptoniphilus_lacrimalis | 0.0434 | | |
| | | Peptoniphilus_lacrimalis | 0.0434 | Fusobacterium_nucleatum | 0.0003 | Fusobacterium_nucleatum | 0.0003 | | |
| | | Fusobacterium_nucleatum | 0.0003 | Clostridiales_bacterium_1_7_47FAA | 0.0342 | Clostridiales_bacterium_1_7_47FAA | 0.0342 | | |
| | | Clostridiales_bacterium_1_7_47FAA | 0.0342 | Porphyromonas_asaccharolytica | 0.0028 | Porphyromonas_asaccharolytica | 0.0028 | | |
| | | Porphyromonas_asaccharolytica | 0.0028 | Porphyromonas_uenonis | 0.0124 | Porphyromonas_uenonis | 0.0124 | | |
| | | Porphyromonas_uenonis | 0.0124 | Peptostreptococcus_stomatis | 0.0076 | Peptostreptococcus_stomatis | 0.0076 | | |
| | | Peptostreptococcus_stomatis | 0.0076 | Campylobacter_ureolyticus | 0.0434 | Campylobacter_ureolyticus | 0.0434 | | |
| | | Campylobacter_ureolyticus | 0.0434 | Clostridium_scindens | 0.0074 | Clostridium_scindens | 0.0074 | | |
| | | Clostridium_scindens | 0.0074 | Parvimonas_micra | 0.0196 | Parvimonas_micra | 0.0196 | | |
| | | Parvimonas_micra | 0.0196 | Dialister_pneumosintes | 0.0010 | Dialister_pneumosintes | 0.0010 | | |
| | | Dialister_pneumosintes | 0.0010 | Firmicutes_bacterium_CAG_534 | 0.0434 | Firmicutes_bacterium_CAG_534 | 0.0434 | | |
| | | Firmicutes_bacterium_CAG_534 | 0.0434 | Bacteroides_pectinophilus | 0.0185 | Bacteroides_pectinophilus | 0.0185 | | |
| | | Bacteroides_pectinophilus | 0.0185 | Streptococcus_gallolyticus | 0.0124 | Streptococcus_gallolyticus | 0.0124 | | |
| | | Streptococcus_gallolyticus | 0.0124 | Methanosphaera_stadtmanae | 0.0231 | Methanosphaera_stadtmanae | 0.0231 | | |
| | | Methanosphaera_stadtmanae | 0.0231 | Megamonas_funiformis_CAG_377 | 0.0434 | Megamonas_funiformis_CAG_377 | 0.0434 | | |
| | | Megamonas_funiformis_CAG_377 | 0.0434 | Peptostreptococcus_anaerobius | 0.0124 | Peptostreptococcus_anaerobius | 0.0124 | | |
| | | Peptostreptococcus_anaerobius | 0.0124 | Eubacterium_infirmum | 0.0434 | Eubacterium_infirmum | 0.0434 | | |
| | | Eubacterium_infirmum | 0.0434 | | | | | | |

| KeohaneDM 2020 – Smoker | | | | | | | | | |
|---|---|---|---|---|---|---|---|---|---|
| **Hyperdimensional Computing** | | **Decision Trees** | | **Logistic Regression** | | **Random Forest** | | **Support Vector Machines** | |
| **Feature** | **p-value** | **Feature** | **p-value** | **Feature** | **p-value** | **Feature** | **p-value** | **Feature** | **p-value** |
| Ruminococcus_sp_CAG_488 | 0.0309 | Holdemania_filiformis | 0.0372 | Ruminococcus_sp_CAG_488 | 0.0309 | Holdemania_filiformis | 0.0372 | Ruminococcus_sp_CAG_488 | 0.0309 |
| Tyzzerella_nexilis | 0.0068 | Ruminococcus_sp_CAG_488 | 0.0309 | Tyzzerella_nexilis | 0.0068 | Ruminococcus_sp_CAG_488 | 0.0309 | Bacteroides_eggerthii | 0.0352 |
| Anaerotruncus_colihominis | 0.0324 | Tyzzerella_nexilis | 0.0068 | Veillonella_dispar | 0.0118 | Tyzzerella_nexilis | 0.0068 | | |
| Streptococcus_australis* | 0.0214 | Veillonella_dispar | 0.0118 | Anaerotruncus_colihominis | 0.0324 | Veillonella_dispar | 0.0118 | | |
| | | Anaerotruncus_colihominis | 0.0324 | Prevotella_bivia | 0.0352 | Anaerotruncus_colihominis | 0.0324 | | |
| | | Prevotella_bivia | 0.0352 | Bacteroides_eggerthii | 0.0352 | Prevotella_bivia | 0.0352 | | |
| | | Bacteroides_eggerthii | 0.0352 | Alistipes_onderdonkii | 0.0168 | Bacteroides_eggerthii | 0.0352 | | |
| | | Alistipes_onderdonkii | 0.0168 | Streptococcus_oralis | 0.0321 | Alistipes_onderdonkii | 0.0168 | | |
| | | Streptococcus_oralis | 0.0321 | Streptococcus_australis | 0.0214 | Streptococcus_oralis | 0.0321 | | |
| | | Streptococcus_australis | 0.0214 | | | Streptococcus_australis | 0.0214 | | |

**KeohaneDM 2020 – Gender**

| Hyperdimensional Computing | | Decision Trees | | Logistic Regression | | Random Forest | | Support Vector Machines | |
|---|---|---|---|---|---|---|---|---|---|
| **Feature** | **p-value** | **Feature** | **p-value** | **Feature** | **p-value** | **Feature** | **p-value** | **Feature** | **p-value** |
| Parabacteroides_merdae | 0.0051 | Parabacteroides_merdae | 0.0051 | Parabacteroides_merdae | 0.0051 | Parabacteroides_merdae | 0.0051 | Parabacteroides_merdae | 0.0051 |
| Parabacteroides_distasonis | 0.0438 | Bacteroides_stercoris | 0.0150 | Agathobaculum_butyriciproducens | 0.0366 | Bacteroides_stercoris | 0.0150 | Bacteroides_stercoris | 0.0150 |
| Eggerthella_lenta | 0.0011 | Agathobaculum_butyriciproducens | 0.0366 | Ruminococcus_bicirculans | 0.0367 | Agathobaculum_butyriciproducens | 0.0366 | Ruminococcus_bicirculans | 0.0367 |
| Holdemania_filiformis | 0.0274 | Parabacteroides_distasonis | 0.0438 | Holdemanella_biformis | 0.0303 | Parabacteroides_distasonis | 0.0438 | Phascolarctobacterium_succinatutens | 0.0064 |
| Holdemanella_biformis | 0.0303 | Odoribacter_splanchnicus | 0.0263 | Phascolarctobacterium_succinatutens | 0.0001 | Odoribacter_splanchnicus | 0.0263 | Ruminococcus_bromii | 0.0023 |
| Phascolarctobacterium_succinatutens | 0.0001 | Eggerthella_lenta | 0.0011 | Ruminococcus_bromii | 0.0023 | Eggerthella_lenta | 0.0011 | Oscillibacter_sp_CAG_241 | 0.0242 |
| Ruminococcus_bromii | 0.0023 | Gordonibacter_pamelaeae | 0.0123 | Slackia_isoflavoniconvertens | 0.0421 | Gordonibacter_pamelaeae | 0.0123 | Eubacterium_siraeum | 0.0066 |
| Slackia_isoflavoniconvertens | 0.0421 | Holdemania_filiformis | 0.0274 | Oscillibacter_sp_CAG_241 | 0.0242 | Holdemania_filiformis | 0.0274 | Mitsuokella_multacida | 0.0427 |
| Oscillibacter_sp_CAG_241 | 0.0242 | Ruminococcus_bicirculans | 0.0367 | Eubacterium_siraeum | 0.0066 | Ruminococcus_bicirculans | 0.0367 | | |
| Eubacterium_siraeum | 0.0066 | Holdemanella_biformis | 0.0303 | Prevotella_sp_AM42_24 | 0.0091 | Holdemanella_biformis | 0.0303 | | |
| Romboutsia_ilealis | 0.0009 | Catenibacterium_mitsuokai | 0.0496 | Prevotella_sp_CAG_1092 | 0.0184 | Catenibacterium_mitsuokai | 0.0496 | | |
| Clostridium_disporicum | 0.0008 | Phascolarctobacterium_succinatutens | 0.0001 | Proteobacteria_bacterium_CAG_139 | 0.0292 | Phascolarctobacterium_succinatutens | 0.0001 | | |
| Brachyspira_sp_CAG_700 | 0.0240 | Slackia_isoflavoniconvertens | 0.0421 | Clostridium_disporicum | 0.0008 | Ruminococcus_bromii | 0.0023 | | |
| Anaerotruncus_sp_CAG_528 | 0.0025 | Oscillibacter_sp_CAG_241 | 0.0242 | Prevotella_sp_885 | 0.0080 | Slackia_isoflavoniconvertens | 0.0421 | | |
| Firmicutes_bacterium_CAG_145 | 0.0075 | Eubacterium_siraeum | 0.0066 | Firmicutes_bacterium_CAG_145 | 0.0075 | Oscillibacter_sp_CAG_241 | 0.0242 | | |
| Actinomyces_turicensis | 0.0256 | Intestinibacter_bartlettii | 0.0156 | Firmicutes_bacterium_CAG_94 | 0.0438 | Eubacterium_siraeum | 0.0066 | | |
| Mitsuokella_multacida | 0.0427 | Clostridium_leptum | 0.0193 | Clostridium_symbiosum | 0.0484 | Intestinibacter_bartlettii | 0.0156 | | |
| | | Prevotella_sp_AM42_24 | 0.0091 | Clostridium_bolteae_CAG_59 | 0.0256 | Clostridium_leptum | 0.0193 | | |
| | | Prevotella_sp_CAG_1092 | 0.0184 | Eisenbergiella_tayi | 0.0484 | Prevotella_sp_AM42_24 | 0.0091 | | |
| | | Proteobacteria_bacterium_CAG_139 | 0.0292 | Streptococcus_sp_F0442 | 0.0240 | Prevotella_sp_CAG_1092 | 0.0184 | | |
| | | Romboutsia_ilealis | 0.0009 | Actinomyces_turicensis | 0.0256 | Proteobacteria_bacterium_CAG_139 | 0.0292 | | |
| | | Clostridium_disporicum | 0.0008 | Mitsuokella_multacida | 0.0427 | Romboutsia_ilealis | 0.0009 | | |
| | | Turicibacter_sanguinis | 0.0006 | | | Clostridium_disporicum | 0.0008 | | |
| | | Prevotella_sp_885 | 0.0080 | | | Turicibacter_sanguinis | 0.0006 | | |
| | | Brachyspira_sp_CAG_700 | 0.0240 | | | Prevotella_sp_885 | 0.0080 | | |
| | | Anaerotruncus_sp_CAG_528 | 0.0025 | | | Brachyspira_sp_CAG_700 | 0.0240 | | |
| | | Firmicutes_bacterium_CAG_145 | 0.0075 | | | Anaerotruncus_sp_CAG_528 | 0.0025 | | |
| | | Firmicutes_bacterium_CAG_94 | 0.0438 | | | Firmicutes_bacterium_CAG_145 | 0.0075 | | |
| | | Alistipes_onderdonkii | 0.0256 | | | Firmicutes_bacterium_CAG_94 | 0.0438 | | |
| | | Clostridium_asparagiforme | 0.0346 | | | Alistipes_onderdonkii | 0.0256 | | |
| | | Clostridium_symbiosum | 0.0484 | | | Clostridium_asparagiforme | 0.0346 | | |
| | | Hungatella_hathewayi | 0.0423 | | | Clostridium_symbiosum | 0.0484 | | |
| | | Clostridium_bolteae_CAG_59 | 0.0256 | | | Hungatella_hathewayi | 0.0423 | | |
| | | Eisenbergiella_tayi | 0.0484 | | | Clostridium_bolteae_CAG_59 | 0.0256 | | |
| | | Streptococcus_sp_F0442 | 0.0240 | | | Eisenbergiella_tayi | 0.0484 | | |
| | | Actinomyces_turicensis | 0.0256 | | | Streptococcus_sp_F0442 | 0.0240 | | |
| | | Mitsuokella_multacida | 0.0427 | | | Actinomyces_turicensis | 0.0256 | | |
| | | | | | | Mitsuokella_multacida | 0.0427 | | |
